# OSGN-1 is a conserved flavin-containing monooxygenase required to stabilize the intercellular bridge in late cytokinesis

**DOI:** 10.1101/2023.08.06.552159

**Authors:** Eugénie Goupil, Léa Lacroix, Jonathan Brière, Sandra Guga, Marc K. Saba-El-Leil, Sylvain Meloche, Jean-Claude Labbé

## Abstract

Cytokinesis is the last step of cell division and is regulated by the small GTPase RhoA. RhoA activity is required for all steps of cytokinesis, including prior to abscission when daughter cells are ultimately physically separated. Like germ cells in all animals, the *C. elegans* embryonic germline founder cell initiates cytokinesis but does not complete abscission, leaving a stable intercellular bridge between the two daughter cells. Here we identify and characterize *C. elegans* OSGN-1 as a novel cytokinetic regulator that promotes RhoA activity during late cytokinesis. Sequence analyses and biochemical reconstitutions reveal that OSGN-1 is a flavin-containing monooxygenase. Genetic analyses indicate that the monooxygenase activity of OSGN-1 is required to maintain active RhoA at the end of cytokinesis in the germline founder cell and to stabilize the intercellular bridge. Deletion of *OSGIN1* in human cells results in an increase in binucleation as a result of cytokinetic furrow regression, and this phenotype can be rescued by expressing a catalytically-active form of *C. elegans* OSGN-1, indicating that OSGN-1 and OSGIN1 are functional orthologs. We propose that OSGN-1 and OSGIN1 are novel, conserved monooxygenase enzymes required to maintain RhoA activity at the intercellular bridge during late cytokinesis and thus promote its stability, enabling proper abscission in human cells and bridge stabilization in *C. elegans* germ cells.

## Introduction

Cell division is a fundamental process that ends with cytokinesis – the physical separation of the two daughter cells. In animal cells, this highly regulated process fundamentally rests on molecules of actin and non-muscle myosin that form an ingressing ring between the two separated sets of sister chromatids at the end of anaphase, eventually forming a transient intercellular bridge that will enable cell abscission (1). The assembly and dynamics of the actomyosin ring are regulated by RhoA, a small GTPase of the Ras superfamily that becomes active upon exchange of its GDP to GTP, mainly by the guanine exchange factor Ect2 (1, 2). Active RhoA coordinates the cortical recruitment and activation of several downstream contractility regulators until its GTP is hydrolyzed into GDP, mostly through the regulated activity of GTPase-activating proteins (GAPs). RhoA is considered as the master regulator of cytokinesis and its activity is thought to be required throughout the process, including in the final stages when it ensures stability of the intercellular bridge and thus favours the proper loading of the abscission machinery.

Cytokinesis does not systematically end with abscission, and incomplete cytokinesis is a conserved feature of germline development in all animals (3-6). This is the case in the embryo of the nematode *C. elegans*, where the germline founder cell, P_4_, undergoes incomplete cytokinesis to give rise to the two primordial germ cells (PGCs), Z_2_ and Z_3_, that remain connected by a stable intercellular bridge (7, 8). Our previous analysis revealed that stability of the PGC intercellular bridge is ensured by a pathway in which RhoA, presumably activated by its upstream regulators Ect2 and centralspindlin, promotes the recruitment of downstream contractility regulators including anillin and non-muscle myosin to stabilize the bridge (7). This work uncovered F30B5.4, an uncharacterized regulator of intercellular bridge stability that genetically clustered with RhoA and its upstream regulators in this pathway (Fig. 1A). Here we characterize F30B5.4 and demonstrate that it codes for a novel, evolutionarily-conserved flavin-dependent monooxygenase enzyme that regulates RhoA activity in late cytokinesis.

**Figure 1.**
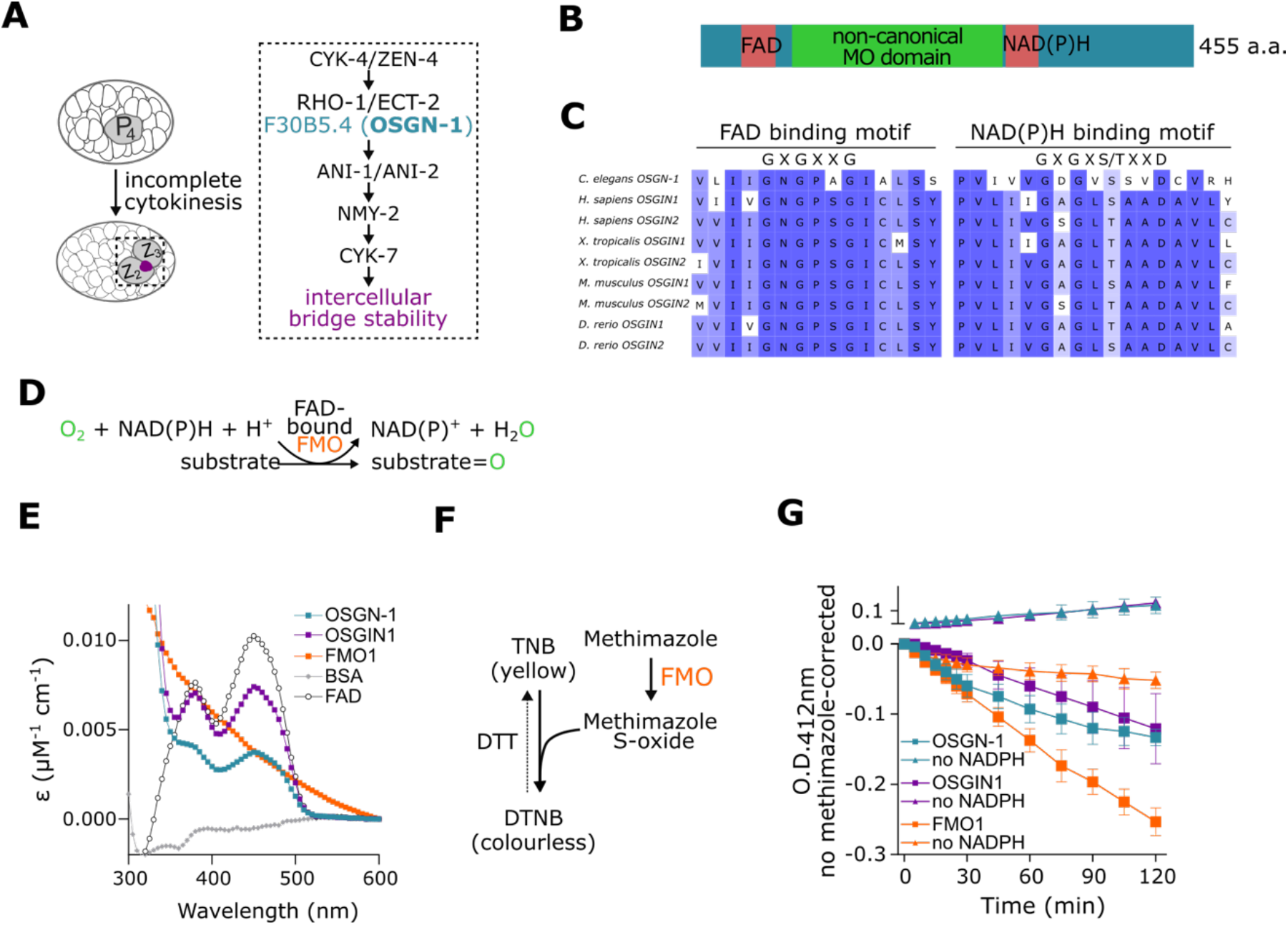
OSGN-1 is a flavin-containing monooxygenase enzyme homologous to human OSGIN1 and OSGIN2. (A) Epistatic pathway mediating intercellular bridge stability following incomplete cytokinesis of the *C. elegans* embryonic germline founder cell, P_4_, into the primordial germ cells (PGCs), Z_2_ and Z_3_. OSGN-1 was previously found to genetically cluster with RHO-1 (RhoA) in this pathway (7). (B) Schematic representation of OSGN-1 organization, showing the putative flavin-adenine dinucleotide (FAD), non-canonical monooxygenase (MO) and nicotinamide adenine dinucleotide phosphate (NADPH) domains. (C) Sequence alignments of the FAD and NADPH domains of *C. elegans* OSGN-1 with those of OSGIN1 and OSGIN2 homologs from different vertebrate species. (D) Depiction of a typical enzymatic reaction catalyzed by flavin-containing monooxygenases (FMO). (E) Extinction coefficient spectra of BSA (negative control, grey), FAD alone (5 μM, white) or FAD bound to human microsomal FMO1 (orange), bacterially-purified *C. elegans* OSGIN1-6xHis (cyan) or human OSGIN1-6xHis (magenta). (F) Schematic depiction of the oxidative reactions in the DTNB-methimazole assay that favour the conversion of two molecules of nitro-5-thiobenzoate (TNB) into one molecule of 5,5’-ditiobis(2-nitrobenzoate) (DTNB). (G) Comparative measurement of the catalytic activity of human microsomal FMO1 (orange), *C. elegans* OSGN-1-6xHis (cyan) and human OSGIN1-6xHis (magenta; 2 μM final concentration each) using the DTNB-methimazole assay, in the presence (squares) or absence (triangles) of the essential co-factor NADPH. The decrease of absorbance at 412 nm over time (in min) reports on catalytic activity. Values are corrected for reactions done in absence of methimazole and correspond to the mean ± SEM over N = 3 assays, see methods for details. Time is relative to addition of methimazole.

## Results and Discussion

### F30B5.4 is a flavin-dependent monooxygenase enzyme homologous to mammalian OSGIN1 and OSGIN2

Comparative sequence analyses revealed that F30B5.4 (hereafter OSGN-1) is conserved across species and is the *C. elegans* homolog of mammalian <u>o</u>xidative <u>s</u>tress-induced <u>g</u>rowth <u>in</u>hibitor 1 and 2 (OSGIN1 and OSGIN2), two proteins that have not yet been extensively characterized. Sequence and structure homology analyses of OSGN-1 showed the presence of two Rossman folds, which are putative binding domains for flavin-adenine dinucleotide (FAD) and nicotinamide adenine dinucleotide (phosphate) (NAD(P)H), flanking a putative, non-canonical monooxygenase (MO) domain (Fig. 1B, C and Fig. S1A). These domains are typically found in so-called flavin-containing monooxygenases (FMOs), a class of enzymes that employ FAD and NAD(P)H as co-factors to catalyze the oxidation of small molecules or protein substrates (Fig. 1D) (9, 10). To test whether OSGN-1 is an FMO, we first assessed the capacity of bacterially-produced and -purified OSGN-1 (fused to 6xHis; Fig. S1B) to bind to FAD *in vitro*. We found that the extinction coefficient profile of OSGN-1 displays the characteristic peaks of FAD absorption (Fig. 1E), consistent with FAD binding (11, 12). To test whether OSGN-1 has monooxygenase activity, we performed the well-established DTNB-methimazole biochemical assay with bacterially-purified OSGN-1 (Fig. 1F) (13-15). We found that OSGN-1 oxidizes methimazole *in vitro*, albeit with different kinetics compared to commercially-produced human FMO1 (Fig. 1G). Interestingly, OSGN-1 was able to directly oxidize TNB into DTNB in this assay (Fig. S1C), an activity that is not seen in FMO1 and that likely impacts its measured activity on methimazole. Importantly however, no activity was measured for OSGN-1 (or FMO1) in absence of the essential co-factor NADPH (Fig. 1G and Fig. S1C), demonstrating that the reaction is catalytic. These results indicate that OSGN-1 is an active FMO.

### OSGN-1 regulates RhoA activity in *C. elegans* PGCs

Our previous analysis suggested that OSGN-1 genetically clusters with the small GTPase RhoA and its upstream regulators in the pathway that promotes maintenance of the PGC intercellular bridge during *C. elegans* embryogenesis (7). This raised the possibility that OSGN-1 regulates RhoA activity in this context. To test this, we employed live imaging of *C. elegans* embryos that express the C-terminal half of the actomyosin scaffold protein anillin fused to GFP (GFP::AHPH), which has previously been shown to robustly report on RhoA activity during cytokinesis in this system (16-19). Monitoring GFP::AHPH during control P_4_ division revealed a peak of fluorescence signal at the cell equator and cytokinetic furrow during anaphase, followed by a maintenance of fluorescence signal at the stable intercellular bridge between the two PGCs (Fig. 2A-C). This spatio-temporal fluorescence pattern is reminiscent of that of actomyosin contractility regulators that ensure PGC intercellular bridge stability downstream of RhoA (7). As reported previously (16), partial RNAi-mediated depletion of RHO-1 (*C. elegans* RhoA (20)) resulted in a decrease of GFP::AHPH cortical fluorescence levels throughout P_4_ division, while depleting the GTPase activating proteins RGA-3/4 (*C. elegans* MP-GAP (21, 22)), which are known redundant signalling terminators of RhoA activity in this context, resulted in an increase of GFP::AHPH fluorescence at the intercellular bridge, compared to control (Fig. 2A-C). These results indicate that the GFP::AHPH reporter behaves as expected in *C. elegans* embryonic germ cells and predictively responds to changes in RhoA activity. Interestingly, RNAi-mediated depletion of OSGN-1 resulted in a decrease of GFP::AHPH cortical fluorescence levels throughout cell division – peak fluorescence levels during anaphase were reduced by half, and the maintenance of fluorescence levels at the stable PGC intercellular bridge in late cytokinesis was abrogated (Fig. 2A-C). These results suggest that OSGN-1 impacts RhoA activity throughout P_4_ cytokinesis and is required for prolonged maintenance of active RhoA at the PGC stable intercellular bridge. Interestingly, while depletion of RHO-1 and RGA-3/4 also impacted GFP::AHPH cortical fluorescence levels at the cytokinetic furrow of dividing somatic cells, the impact of OSGN-1 depletion was only observed in P_4_ (Fig. S2), indicating that its effect on RhoA activity is restricted to the germ lineage. This is compatible with published RNA-seq data in which *osgn-1* mRNA levels were found to be below detection levels in most early embryonic blastomeres and become enriched specifically in the P_4_ blastomere (23).

**Figure 2.**
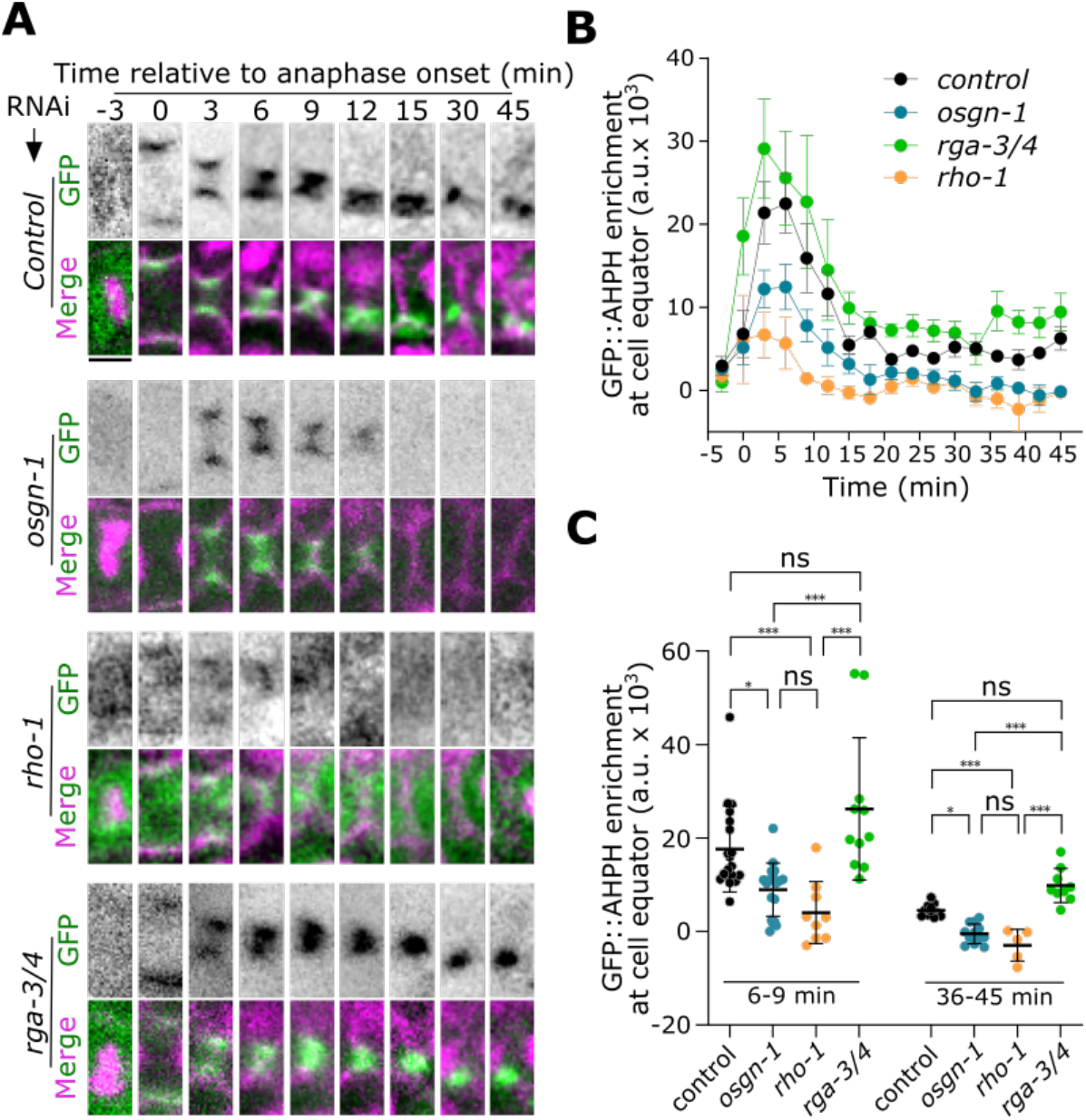
OSGN-1 regulates RhoA activity at the *C. elegans* PGC intercellular bridge. (A) Time-lapse images (sum projections of three confocal slices) of the germline founder cell division in *C. elegans* embryos depleted of control (empty L4440 vector), *osgn-1*, *rho-1* or *rga-3/4* by RNAi. Embryos express markers for RhoA activity (GFP::AHPH, green), membrane (mCh::PLC∂-PH, magenta) and chromatin (mCh::H2B, magenta). Time (in min) is relative to anaphase onset (A.O.). Scale bar = 3 μm. (B) Fluorescence levels over time of the GFP::AHPH reporter measured at the equatorial region (cytokinetic furrow/intercellular bridge) of P_4_ in embryos depleted of control (black), *osgn-1* (blue), *rho-1* (orange) or *rga-3/4* (green) by RNAi, as in panel a. Values correspond to mean ± SEM over 3 biological replicates (n=5-19 embryos in total). (C) Average fluorescence intensity of the GFP::AHPH reporter measured during P_4_ cytokinetic furrow ingression (6-9 min after A.O.) and PGC intercellular bridge maintenance (36-45 min after A.O.) in embryos depleted of the indicated gene products. Values represent individual embryos. Bars denote average ± SD (n=5-19 embryos). ns = non-significant, *p < 0.05, ***p < 0.001, two-way ANOVA followed by a Šídák multiple comparison test.

### The monooxygenase activity of OSGN-1 is required for its function at the PGC intercellular bridge but is dispensable for fertility

To assess whether the monooxygenase activity of OSGN-1 is required for its function we phenotypically characterized 13 available *C. elegans* strains bearing point mutations across the *osgn-1* gene (24). The mutations introduce amino acid substitutions in all three domains of OSGN-1 (FAD, NADPH, MO) and two of them introduce a premature stop codon, one of these (W^201^Stop) being predicted to truncate the protein in the middle of the MO domain (Fig. 3A and Fig. S3A). We found that 11 of these mutations resulted in a significant proportion of embryos in which the PGC intercellular bridge was unstable and eventually regressed, prior to the start of epidermal morphogenesis (Fig. 3B, C and Fig. S3B). This defect is similar to the binucleated PGC phenotype observed following RNAi-mediated depletion of OSGN-1 or actomyosin contractility regulators (7). Surprisingly, all mutations had only a marginal impact on fertility, brood size and embryonic viability, independently of the degree to which they perturbed intercellular bridge stability (Fig. S3B-E). Extended time-lapse analysis of embryos from 3 of these mutant strains revealed that most PGC membrane partitions that had regressed eventually reformed and ended up with seemingly mononucleated PGCs, as is observed in control embryos (Fig. S3F, G). This indicates the existence of a mechanism, later during embryonic development, that compensates for the loss of intercellular bridge stability in PGCs to ensure proper partitioning between these cells. Interestingly, the membrane partition rescue occurs around the time when PGCs are reported to initiate the formation of polar lobes (25, 26), raising the possibility that the mechanism enabling polar lobe formation can compensate for the destabilization of the PGC intercellular bridge.

**Figure 3.**
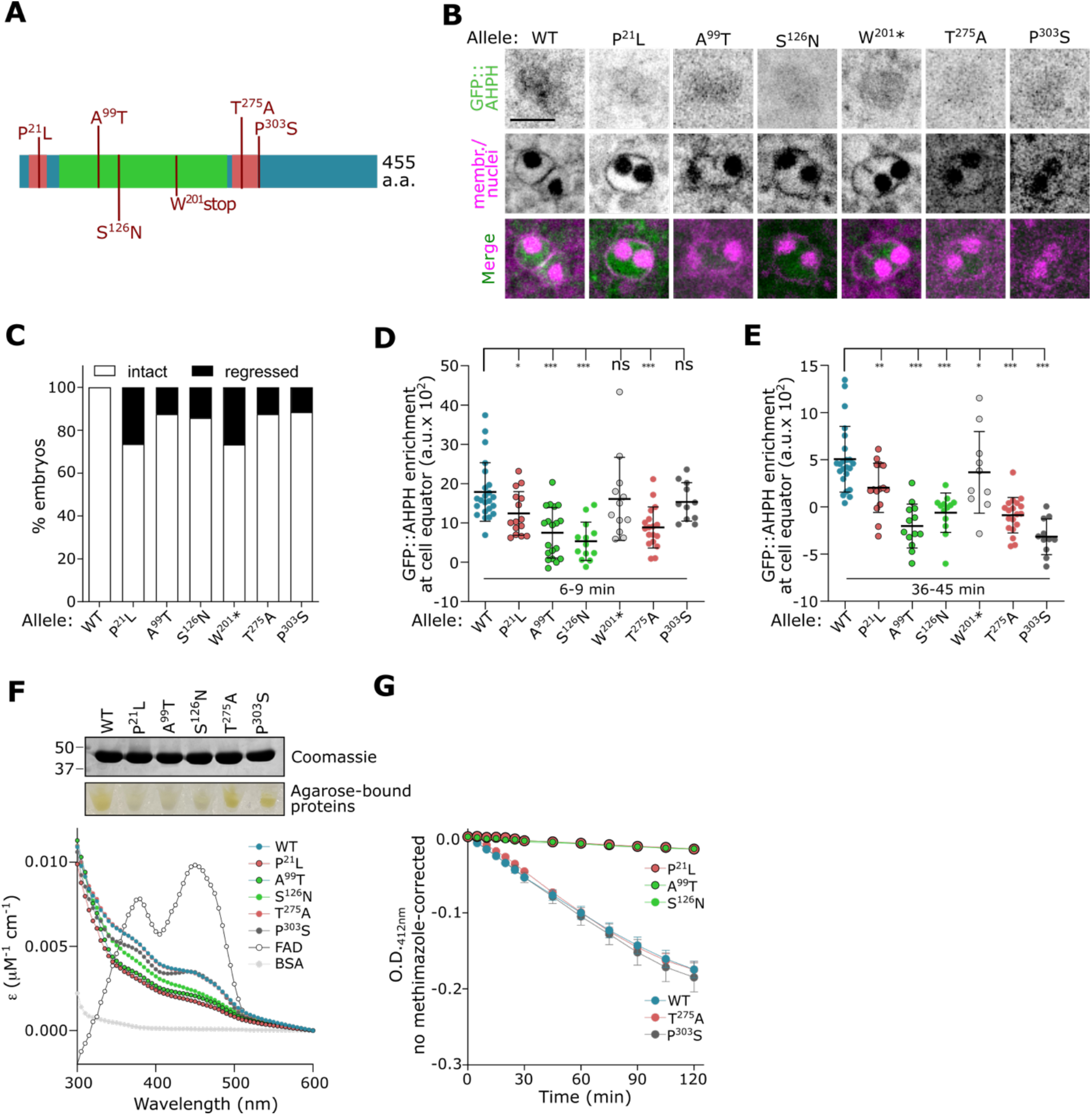
The catalytic activity of OSGN-1 is required for stability of the *C. elegans* PGC intercellular bridge. (A) Schematic depiction of the different amino acid substitutions encoded by some of the *osgn-1* mutants used in this study. (B-C) Representative images (B) and quantification (C) of the presence (white bars) or absence (black bars) of a membrane partition between the two PGCs in embryos expressing markers for RhoA activity (GFP::AHPH, green), membrane (mCh::PLC∂-PH, magenta) and chromatin (mCh::H2B, magenta). Scale bar = 10 μm, n = 16-57 embryos. (D-E) Average fluorescence intensity of the GFP::AHPH reporter measured during P_4_ cytokinetic furrow ingression (6-9 min after anaphase onset [A.O.]; D) and PGC intercellular bridge maintenance (36-45 min after A.O.; E) in embryos encoding the indicated OSGN-1 variants. Values represent individual embryos. Bars denote average ± SD (n=13-23 embryos). Individual fluorescence curves are shown in Fig. s3h. ns = non-significant, *p < 0.05, **p < 0.01, ***p < 0.001, one-way ANOVA followed by a Dunnett’s multiple comparison test. (F) Extinction coefficient spectra of BSA (negative control, grey), FAD alone (5 μM, white) or FAD bound to the bacterially-purified OSGN-1 variants indicated (WT and mutants, colored). Top inset, Coomassie staining of purified OSGN-1 variants used in the FAD-binding assay. Bottom inset, Ni-NTA agarose pellets following OSGN-1 variant purifications, visually denoting the presence (yellowish coloration) or absence (white) of FAD. (G) Comparative measurement of the catalytic activity of the indicated *C. elegans* OSGN-1-6xHis variants (1 μM final concentration each) using the DTNB-methimazole assay, in the presence of the essential co-factor NADPH. The decrease of absorbance at 412 nm over time (in min) reports on catalytic activity. Values are corrected for reactions done in absence of methimazole and correspond to the mean ± SEM over N = 3 assays, see methods for details.

We next sought to determine the impact of OSGN-1 amino acid substitutions on RhoA activity at the PGC stable intercellular bridge *in vivo* using the GFP::AHPH reporter, and on OSGN-1 catalytic activity *in vitro* using the DTNB-methimazole assay. This was done with a subset of mutants that differentially impacted *osgn-1*(lf)-associated phenotypes, namely intercellular bridge stability (P^21^L, W^201^Stop), fertility (S^126^N, P^303^S), embryonic viability (A^99^T) and brood size (T^275^A; Fig. 3A). The fluorescence intensity of the GFP::AHPH reporter was scored in these mutants at peak intensity during anaphase and during signal persistence after telophase, highlighting the relative activity of RhoA during cytokinetic furrow ingression and maintenance of the PGC intercellular bridge, respectively. We found that all mutations tested decreased GFP::AHPH cortical fluorescence intensity during peak and maintenance phases except W^201^Stop and P^303^S, which impacted reporter fluorescence levels during maintenance but had no effect on peak recruitment in early cytokinesis (Fig. 3D, E and Fig. S3H). Using the DTNB-methimazole assay, we found that bacterially-purified OSGN-1 variants bearing the P^21^L, A^99^T and S^126^N amino acid substitutions did not bind to FAD (Fig. 3F) and had no measurable monooxygenase activity *in vitro* (Fig. 3G). Interestingly, the T^275^A and P^303^S C-terminal variants of OSGN-1 were as active as control in these *in vitro* biochemical assays. As animals bearing these non-conservative amino acid substitutions display phenotypic defects comparable to those observed in mutants producing catalytically-inactive OSGN-1, this indicates that these mutations impact a functionally-relevant activity of OSGN-1 that is independent of its monooxygenase activity. Together, these results indicate that while the loss of OSGN-1 catalytic activity has variable impacts on fertility, brood size, embryonic viability and peak intensity of the GFP::AHPH reporter at the onset of cytokinetic furrow ingression, it consistently results in a strong decrease of PGC intercellular bridge stability and GFP::AHPH reporter levels at the stable bridge. We conclude that the monooxygenase activity of OSGN-1 regulates RhoA activity throughout P_4_ cytokinesis but is mainly required after furrow ingression to promote the maintenance of the PGC intercellular bridge stability.

### The function of OSGN-1 in cytokinesis is conserved in human cells

Our results raised the possibility that OSGN-1 functions at the PGC stable intercellular bridge to locally regulate RhoA activity. We sought to test this directly in *C. elegans* by engineering animals endogenously expressing OSGN-1 fused to mNeonGreen (using CRISPR/Cas9-mediated gene editing), but expression levels were too low to enable detection, and polyclonal antibodies raised against OSGN-1 also failed to reveal a specific signal in fixed *C. elegans* embryos. Therefore, to determine whether OSGN-1 can localize to regions where RhoA is typically active during M phase, we instead employed HeLa cells as heterologous system in which we expressed a human codon-optimized form of *C. elegans* OSGN-1 fused to GFP (GFP-HA-OSGN-1; Fig. 4A) under the control of the constitutive cytomegalovirus promoter. Using confocal time-lapse imaging and immunofluorescence of HeLa cells undergoing mitosis, we found that OSGN-1 was detected diffusely throughout the cytoplasm until late telophase, when it became enriched at the midbody and colocalized with RhoA and pMLC (Fig. 4B, C and Fig. S4A). These results indicate that *C. elegans* OSGN-1 can be recruited to the midbody in dividing HeLa cells.

**Figure 4.**
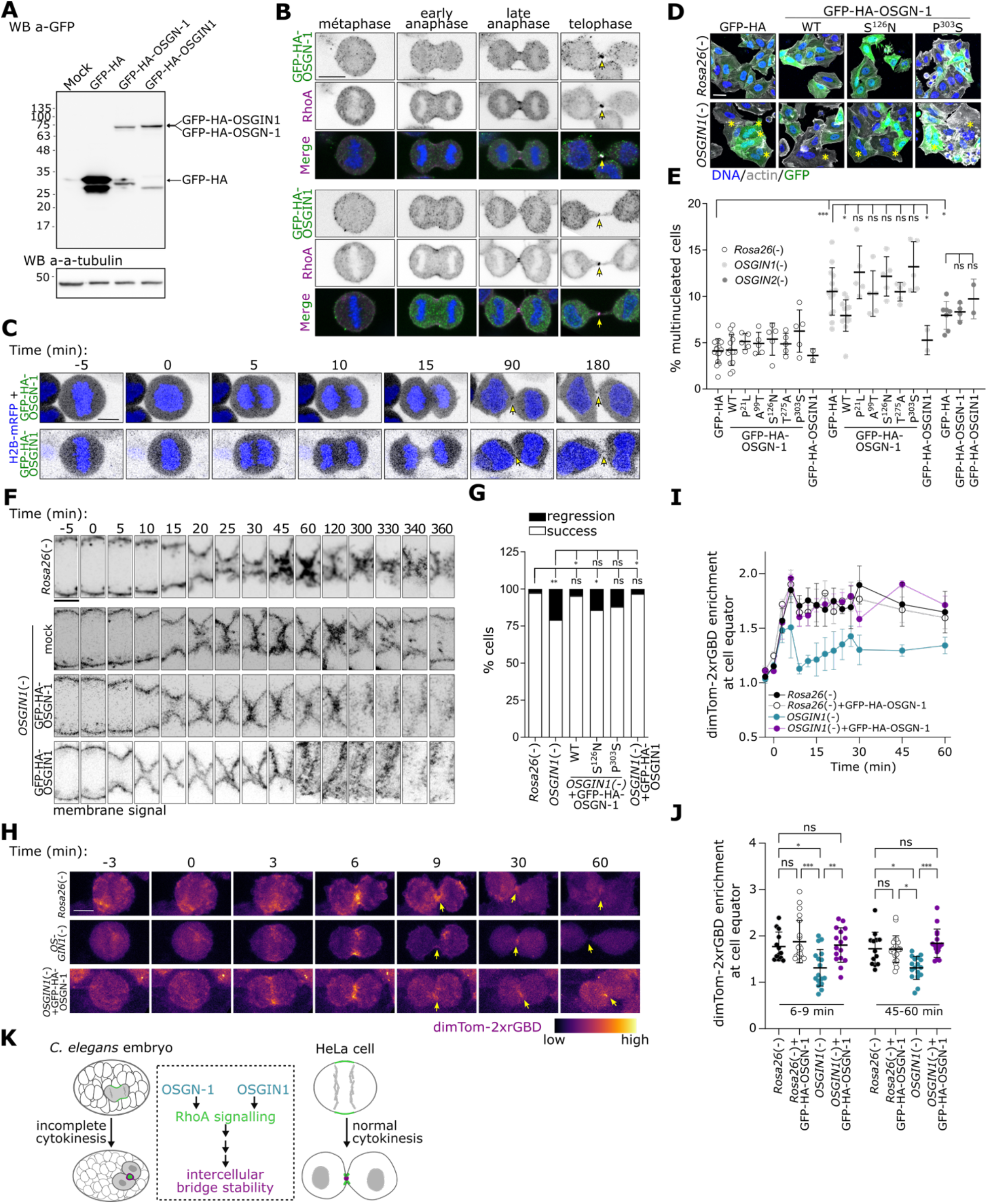
The function of OSGN-1 in cytokinesis is conserved in human cells. (A) Western blot analysis (revealed with anti-GFP antibodies) of extracts from HeLa cells untransfected (mock) or expressing either GFP-HA, GFP-HA-OSGN-1 or GFP-HA-OSGIN1. (B) Representative indirect immunofluorescence images of HeLa cells stably expressing GFP-HA-OSGN-1 (top set) or GFP-HA-OSGIN1 (bottom set) and stained with antibodies against GFP (green) and RhoA (magenta) at the indicated mitotic stages. Hoechst labels nuclei (blue). Arrows denote the intercellular bridge. Scale bar = 10 μm. (C) Time-lapse images of HeLa cells stably expressing H2B-mRFP (blue) and GFP-HA-OSGN-1 (top row) or GFP-HA-OSGIN1 (bottom row). Arrows denote the intercellular bridge. Time (in min) is relative to anaphase onset. Scale bar = 10 μm. (D) Representative images of HeLa cells deleted of *Rosa26* (control) or *OSGIN1*, and transiently expressing either GFP-HA or the indicated GFP-HA-OSGN-1 variants (green). Asterisks denote multinucleated cells. Hoechst labels nuclei (blue) and phalloidin labels F-actin (white). (E) Quantification of the degree of multinucleation in HeLa cells deleted of *Rosa26* (control), *OSGIN1* or *OSGIN2*, and stably expressing either GFP-HA, the indicated GFP-HA-OSGN-1 variants or GFP-HA-OSGIN1. ns = non-significant, *p < 0.05, **p < 0.01, ***p < 0.001, two-way ANOVA followed by a Tukey’s multiple comparison test, in N>3 biological replicates (n>720 cells in total). (F-G) Time-lapse images of cytokinetic furrow ingression (F) and quantification of cytokinesis outcome (G) in *Rosa26* control or *OSGIN1* mutant HeLa cells that stably express markers for the plasma membrane (FP-CAAX) and mock (second row) GFP-HA-OSGN-1 (third row) or GFP-HA-OSGIN1 (bottom row; see also Fig. s4d). Time is relative to anaphase onset. Scale bar = 10 μm. ns = non-significant, *p < 0.05, **p < 0.01, Fisher’s exact test, in N > 3 biological replicates (n=28-74 cells). (H-I) Representative images (H) and fluorescence levels over time (I) of the dimericTomato-2xrGBD RhoA activity reporter (colored scale) measured at the equatorial region (cytokinetic furrow/intercellular bridge) of HeLa cells deleted of *Rosa26* (control) or *OSGIN1*, either mock transfected (top and middle panels) or co-expressing (bottom panel) GFP-HA-OSGN-1. Time is relative to anaphase onset. Arrows denote the intercellular bridge. Scale bar = 10 μm. Values correspond to mean ± SEM over N > 3 biological replicates (n=13-21 cells in total). (J) Average fluorescence intensity of the dimericTomato-2xrGBD reporter measured in early (6-9 min after anaphase onset) or late (45-60 min after anaphase onset) HeLa cell cytokinesis. Values represent individual cells and bars denote average ± SD (n=13-21 cells). ns = non-significant, *p < 0.05, **p < 0.01, ***p < 0.001 two-way ANOVA followed by a Tukey multiple comparison test. (K) Proposed model for the regulation of cytokinesis by OSGN-1 and OSGIN1 proteins.

Public datasets and our qPCR analyses (Fig. S4B) indicate that HeLa cells express both *OSGIN1* and *OSGIN2*, the two human homologs of *C. elegans osgn-1*. To test if the role of OSGN-1 in cytokinesis is conserved, we first assessed whether CRISPR/Cas9-mediated deletion of either human *OSGIN* gene impacts cell division. We found that HeLa cells lacking *OSGIN1* display a significant increase in the proportion of multinucleated cells (2 or more nuclei) compared to *Rosa26* mutant control (Fig. 4D, E). Time-lapse analysis of dividing cells revealed that mitotic duration and the rates of furrow ingression in *OSGIN1* mutant cells are comparable to *Rosa26* control (Fig. S4C), but that the membrane regresses in a proportion of cells at the end of cytokinesis, resulting in binucleation (Fig. 4F, G and Fig. S4D; Movie 1). Furthermore, monitoring of a RhoA activity reporter (the Rho-binding domain of Rhotekin fused to dTomato (27)) during HeLa cell cytokinesis suggested a decrease of RhoA activity at the cytokinetic furrow and intercellular bridge in cells lacking *OSGIN1* compared to *Rosa26* control cells (Fig. 4H-J; Movie 2). These defects are reminiscent of the loss of *osgn-1* in *C. elegans* germ cells and suggest that human OSGIN1 impacts HeLa cell ploidy by promoting RhoA activity during cytokinesis and thus ensuring cytokinetic completion. Strikingly, expression of wild-type *C. elegans* OSGN-1 in HeLa cells lacking *OSGIN1* completely rescued these defects and restored ploidy (Fig. 4D, E), membrane stability in late cytokinesis (Fig. 4F, G and and Fig. S4D) and fluorescence levels of the RhoA activity reporter (Fig. 4H-J), all to proportions comparable to control. Importantly, this rescue by *C. elegans* OSGN-1 required its MO activity, as expressing the catalytically-inactive OSGN-1^P21L^, OSGN-1^A99T^ or OSGN-1^S126N^ variants did not rescue ploidy defects (Fig. 4D, E) or membrane stability (Fig. 4F, G and Fig. S4D). Furthermore, no rescue was observed after expression of the catalytically-active but otherwise non-functional OSGN-1^T275A^ or OSGN-1^P303S^ variants (Fig. 4D-G and Fig. S4D), indicating that the MO-independent activity of OSGN-1 is also needed in this context. CRISPR/Cas9-mediated deletion of *OSGIN2* also led to an increase in the proportion of multinucleated HeLa cells compared to control, but this phenotype was not rescued by expression of *C. elegans* OSGN-1 (Fig 4E). These results indicate that *C. elegans* OSGN-1 is functionally orthologous to human OSGIN1 and further highlight the requirement of the MO activity of OSGN-1 to mediate its function.

### Human OSGIN1 is a flavin-containing monooxygenase

The phenotypic rescue of *OSGIN1* mutant HeLa cells by *C. elegans* OSGN-1 suggested that human OSGIN1 is also an FMO. To test this, we assessed the activity of bacterially-purified OSGIN1 (fused to 6xHis; Fig S1B) using the DTNB-methimazole biochemical assay. We found that human OSGIN1 binds to FAD and oxidizes methimazole *in vitro* in a manner comparable to *C. elegans* OSGN-1 (Fig. 1E, G and Fig S1C), demonstrating that it is an active FMO. Expressing OSGIN1 (fused to GFP-HA) in HeLa cells revealed that it localizes to the midbody in late cytokinesis (Fig. 4B, C) and that it rescues the binucleation (Fig. 4E) and membrane regression (Fig. 4F, G and Fig. S4D) defects of *OSGIN1* mutant cells. Furthermore, overexpression of OSGIN1 in *OSGIN2* mutants did not rescue multinucleation defects (Fig. 4E). These results indicate that, as is the case for *C. elegans* OSGN-1, human OSGIN1 is an active FMO that localizes to the midbody and impacts intercellular bridge stability in late cytokinesis.

Taken together, our work demonstrates that OSGN-1 and OSGIN1 are novel cytokinetic regulators that are recruited at the midbody in late cytokinesis to locally promote RhoA activity. In *C. elegans* germline founder cells that undergo incomplete cytokinesis, this contributes to maintain the stability of the intercellular bridge until the onset of epidermal morphogenesis. In HeLa cells, this contributes to transiently stabilize the intercellular bridge while cells prepare for abscission (Fig. 4K). The finding that both OSGN-1 and OSGIN1 proteins are active FMOs and that expression of *C. elegans* OSGN-1 can rescue the deletion of *OSGIN1* in HeLa cells indicates that these proteins are functional orthologs. Furthermore, the lack of phenotypic rescue of *OSGIN2* mutant HeLa cells by expression of either *C. elegans* OSGN-1 or human OSGIN1 suggests that the two human OSGIN proteins are not functionally redundant. Previous studies demonstrated that MICAL, another flavin-containing monooxygenase, promotes actin oxidation at the midbody to facilitate abscission (28, 29). While the substrate of OSGN-1/OSGIN1 is currently unknown, these proteins do not belong to the same class of FMOs as MICAL and are thus unlikely to carry out their activity via actin oxidation. Nonetheless, both types of monooxygenases have a significant impact on abscission, suggesting that catalytic oxidation may be a prevalent means to fine tune cellular activities that underpin key cellular processes.

## Materials and Methods

### Sequence analyses and structural comparisons

The amino acid sequence of *C. elegans* F30B5.4a (WormBase WS265) was analyzed using BLASTP against non-redundant sequences in the UniProtKB/SwissProt databases (30), and *H. sapiens* OSGIN1 and OSGIN2 were the homologs found with highest similarity. Alignments with OSGIN1 and OSGIN2 proteins from different species were done using the T-COFFEE algorithm of Ugene software (31, 32). For fold homology analysis, the I-TASSER algorithm (33, 34) was used to generate a theoretical 3D structure, and the templates used to generate this theoretical structure suggested that F30B5.4a (OSGN-1) is a flavin-containing monooxygenase.

### Protein purification and DTNB-methimazole assay

The coding sequence of *C. elegans osgn-1* (F30B5.4a, WormBase WS265) and human *OSGIN1* were amplified by PCR using Phusion DNA polymerase (NEB, M0530L), according to manufacturer’s instructions, cloned into bacterial expression vector pET-22 to insert a 6xHis tag at its C-terminal (OSGN-1-6xHis and OSGIN1-6xHis, see section on DNA constructs below for more details) and transformed in *E. coli* strain BL21(DE3). Bacteria were grown in LB broth at 37°C to an O.D. of ∼0.7-0.8 and induced with 1 mM IPTG in the presence of 0.05 mM riboflavin (Sigma, R9504) overnight at 18°C. All subsequent steps were performed on ice. Cells were collected by centrifugation, resuspended in lysis buffer (50 mM Tris pH 8.0, 150 mM NaCl, 20 mM imidazole, 1mM DTT, 1 mM PMSF, 0.1 % Triton X-100) using 1/10 of the volume of the culture medium and sonicated 6 x 10 sec (at 50% intensity, using a Fisher Scientific FB120 sonicator). Lysates were spun at 18 000 x g to remove cell debris, and incubated with Nickel-NTA agarose (QIAGEN, 1018244) for 2h at 4°C. Beads were washed twice in ten bed volumes of lysis buffer and twice in ten bed volumes of HBS buffer (20 mM HEPES pH 7.5, 150 mM NaCl, 2 mM MgCl_2_, 20 mM imidazole, 1 mM DTT). Bound material was eluted from the agarose matrix by incubating in 1 ml elution buffer (50 mM Tris pH 8.0, 500 mM NaCl, 5 mM MgCl_2_, 300 mM imidazole), twice 15 min at 4°C. Eluates were passed through a Zeba spin desalting column (7k MWCO, Thermo Scientific, PI89892) saturated with storage buffer (50 mM sodium phosphate buffer pH 7.4, 150 mM NaCl, 10% glycerol), aliquoted and flash frozen in liquid nitrogen for further use. Protein purity was verified by SDS-PAGE, followed by Coomassie staining. FAD absorption spectra were conducted on 5-10 µM of freshly-purified protein in PBS buffer, by measuring absorption between 260 nm and 550 nm (with 5 nm increments) using a Tecan Infinite M200 Pro reader (Tecan Life Sciences), as previously (11, 12). Variants of OSGN-1 bearing single amino acid substitutions were expressed and purified using the same method.

The DTNB-methimazole assay was performed with technical duplicates in 96-well plates (Greiner, 655201), as described previously (13, 14). Reactions were conducted in a final volume of 300 µl of Tricine/KOH buffer (100 mM Tricine pH 8.5, 2.5 mM EDTA) supplemented with 0.1 mM nicotinamide adenine phosphodinucleotide phosphate (NADPH, Sigma, 10107824001), 60 nM 5,5′-dithiobis(2-nitrobenzoic acid) (DTNB, Sigma, D8130) and 120 nM 1,4-Dithiothreitol (DTT, Sigma). Commercially-available human FMO1 (Sigma, F4928-1VL), bacterially-purified human OSGIN1-6xHis or bacterially-purified *C. elegans* OSGN-1-6xHis (wild-type and variants) were added to a final concentration of 1-2 µM. Reactions were started by addition of methimazole (Sigma, 46429) to a final concentration of 1 mM (from a fresh 300 mM stock solution in methanol) and monitored for 3h at room temperature by optical densitometry at 412 nm using a Tecan Infinite M200 Pro reader (Tecan Life Sciences). Raw optical densities for each reaction were first normalized by subtracting the value at time zero (upon addition of methimazole or methanol control), and datasets conducted in the presence of methimazole were further normalized by subtracting at each time point the corresponding value in the methanol control condition (no methimazole), as done previously (13, 14).

### *C. elegans* strain maintenance, genetics and RNAi depletions

Animals were cultured as previously described (35) and grown on nematode growth media (NGM) at 20°C. For all experiments, worms were synchronized at the L1 larval stage by hatching embryos in M9 buffer (0.022 mM KH_2_PO_4_, 0.042 mM Na_2_HPO_4_, 0.086 mM NaCl, 1 mM MgSO_4_) after sodium hypochlorite treatment (1.2% NaOCl, 250 mM KOH), and allowed to grow on OP50 bacteria to the desired developmental stage. Brood size was measured by transferring 10 L4-stage animals of a given genotype to individual NGM/OP50 plates and allowing them to lay eggs for 5 days at 20°C, with a daily transfer to a new plate (to facilitate counting), and manually counting the total number of eggs laid per animal. Embryonic viability was assessed 24h later by scoring the percentage of these eggs that hatched into L1 larvae. Fertility was assessed by scoring the percentage of hatched progeny that accumulated embryos in their uterus upon reaching adulthood. The strains used in this study are listed in Table S1.

Strains bearing *osgn-1* mutant alleles were generated through the Million Mutation Project (24) and obtained from the *Caenorhabditis* Genetics Center (University of Minnesota). Each strain was outcrossed a minimum of 10 times using strain UM729, containing the *dpy-13(e184)* mutation that served as a visible marker in *trans* of the *osgn-1* locus. The presence of each *osgn-1* allele in the final homozygote strains was confirmed by Sanger sequencing (see Table S2 for oligonucleotide sequences). All strains contained markers enabling the visualization of active RhoA (GFP::AHPH (16)), chromatin (mCherry::HIS-58 (36)) and plasma membrane (mCherry::PH^PLC61^ (37)).

RNAi depletions were done by soaking animals in dsRNA solution, as described previously (38). Briefly, plasmids containing sequences of interest inserted between two T7 promoters in the L4440 vector (*rho-1*: cenix:169-h12; *osgn-1*: sjj_F30B5.4; *rga-3/4*: sjj_K09H11.3) were subjected to PCR amplification with T7-specific oligonucleotides and Phusion DNA polymerase. The PCR products were cleaned with QIAquick® gel extraction kit (Qiagen, 28706) and used as template to generate dsRNA *in vitro* using the Promega T7 Ribomax^TM^ Express RNAi System kit (Promega, P1320). The dsRNA was precipitated with sodium acetate (300 mM final) and an equal volume of isopropanol, washed with ethanol and resuspended in nuclease-free water. Three to five synchronized L4 larvae were rinsed in M9 buffer and individually transferred to a tube containing 5 µl of RNAi soaking buffer (10.9 mM Na_2_HPO_4_, 5.5 mM KH_2_PO_4_, 2.1 mM NaCl, 4.7 mM NH_4_Cl, 6.3 mM spermidine, 0.11% gelatin) supplemented with dsRNA targeting *rho-1* (0.5 µg), *osgn-1* (5 µg) or *rga-3/4* (1 µg). After 24h incubation at 20°C, animals were recovered for 48h on NGM/OP50 plates prior to assays.

### DNA constructs

The oligonucleotide and gene block sequences used in this study are available in Table S2, and the vectors are detailed in Table S3. For all constructs detailed below, PCR amplifications were done using Phusion DNA polymerase (NEB, M0530L), following manufacturer’s instructions, and sequences were verified by Sanger sequencing.

To generate the pCMV-H2B-mRFP1 construct enabling visualization of H2B-mRFP in HeLa cell nuclei, the H2B-mRFP1 sequence was amplified by PCR from pLV-RFP (Addgene #26001 (39)) and inserted using Gibson assembly in place of GFP into vector pEGFP-C1 (Clonetech, a gift from S. Carréno).

To generate the pcDNA3.1(+)-mRFP1-CAAX construct enabling the targeting of mRFP at the membrane of HeLa cells, mRFP1 was amplified by PCR from pCMV-H2B-mRFP1 and inserted using Gibson assembly in place of GFP into vector pcDNA3.1(+)-rGFP-CAAX (40).

To generate the pEGFP-HA-OSGN-1 or pEGFP-HA-OSGIN1 constructs, we first used PCR to introduce a hemagglutinin (HA) tag C-terminal of GFP into vector pEGFP-C1, thus generating the pEGFP-HA-C1 control vector. We then employed gene synthesis services (Integrated DNA Technologies) to obtain a human codon-optimized version of the *osgn-1* cDNA (hereafter *HsF30B5.4a*, see Table S3 for complete sequence) or human *OSGIN1* cDNA (with some silent codon alterations from the canonical *OSGIN1* sequence, see Table S3), amplified it by PCR adding EcoRI and BamHI (OSGN-1) or SacI and KpnI (OSGIN1) restriction sites on each end, and used these sites to insert this fragment into pEGFP-HA-C1.

To generate pRK5-HA-OSGN-1 or pRK5-HA-OSGIN1 constructs, the HA-OSGN-1 or HA-OSGIN1 sequences were amplified by PCR from the pEGFP-HA-OSGN-1 or pEGFP-HA-OSGIN1 constructs, with ClaI and BamHI (OSGN-1) or ClaI and SalI (OSGIN1) restriction sites on each end, and inserted at these sites into vector pRK5 (BD Pharmigen, a gift from Audrey Claing). To generate the pET-22b-OSGN-1 and pET-22b-OSGIN1 constructs, enabling production of OSGN-1-6xHis and OSGIN1-6xHis in bacteria, the *HsF30B5.4a* or *OSGIN1* sequences were amplified by PCR from the pEGFP-HA-OSGN-1 or pEGFP-HA-OSGIN1 constructs, adding NdeI and XhoI restriction sites on each end, and inserted at these sites into vector pET-22b (Novagen, a gift from Mike Tyers). The generation of constructs enabling expression of OSGN-1 variants in bacteria and HeLa cells was done by site-directed mutagenesis of the pET-22b-OSGN-1 and pEGFP-HA-OSGN-1 vectors, respectively, using the same oligonucleotides to introduce each variant in both constructs. Oligonucleotides introducing a mutation at a specific site for each variant (see Table S3) were used in PCR reactions on these vectors, as done previously (41). PCR products for each mutagenic pair were then denatured by heat and mixed into a single tube, digested with DpnI to degrade template strands, and transformed into DH5a competent *E. coli* cells for colony screening and clone recovery.

The single guide (sg) RNA sequences used to target *Rosa26*, *OSGIN1* and *OSGIN2* were synthesized (Integrated DNA Technologies; see Table S2 for sequences) and cloned as oligonucleotide dimers into BsmBI-digested plentiCRISPR-v2-puro (Addgene #98290, a gift from Feng Zhang), as described previously (42).

### Human cell culture, transfection and lentiviral transduction

Human cell lines were grown at 37°C with 5% CO_2_ in Dulbecco’s Modified Eagle’s Medium (DMEM, Sigma, D6429) supplemented with 10% (v/v) heat-inactivated fetal bovine serum (Cytiva, SH30396.03), 100 U/ml penicillin and 100 μg/ml streptomycin (Invitrogen, 15140122). For transient transfection, cells seeded at a density of 1 × 10^6^ cells per 10 cm dish, 1.5-3 × 10^5^ cells per well in a 6-well plate, or 1.5-3 × 10^5^ cells per 35 mm microscopy plate, were transfected using Lipofectamine 3000 treatment (Invitrogen, L3000015) according to manufacturer’s instructions. Experiments were performed 36-48h post-transfection. Stable cell lines expressing GFP-HA alone, or co-expressing GFP-HA-OSGIN1 or GFP-HA-OSGN-1 (wild-type and variants) were generated by transfection (as above) and subsequent selection for 14 days in growth medium supplemented with 1.5 mg/mL G418 sulfate solution (Wisent, 450-130-QL).

The generation of HeLa cells knocked out for *Rosa26*, *OSGIN1* or *OSGIN2* was done by CRISPR/Cas9-mediated gene editing, as described previously (42, 43). HEK293T cells (1.5 x 10_6_) were first seeded in a 10 cm dish and grown for 24h. The following day, cells were co-transfected (using Lipofectamine 3000 treatment, as described above) with pMD2.g (envelope plasmid, Addgene #12259), psPAX2 (packaging plasmid, Addgene #12260, a gift from Didier Trono) and lentiCRISPRv2-puro (transfer plasmid) containing a small guide (sg) RNA for either human *OSGIN1* (sg1 to sg5), *OSGIN2* (sg1 to sg6) or *Rosa26* (sg1; see Table S2 for sequences). The following day, the medium was changed with fresh medium containing 20% FBS and the cells were incubated for an additional 36h. The medium of each culture was then harvested, centrifuged for 5 min at 860g to remove debris, and filtered using a 0.45 µm surfactant-free cellulose acetate syringe filter (Corning, 431220). To transduce HeLa cells, this filtered lentivirus-containing medium was applied directly to HeLa cells that had been plated at a density of 1 x 10^6^ cells in a 10 cm dish 24h before, and hexadimethrine bromide (Sigma 107689) was added to a final concentration of 8 µg/ml. The next day, the medium was changed for medium containing 1 µg/mL puromycin for selection of transduced cells. HeLa cells mutant for *Rosa26* were kept as a polyclonal population. For *OSGIN1*, the efficiency of each sgRNA (sg1 to sg5) was verified by extracting genomic DNA (Monarch Genomic DNA Purification Kit, NEB T3010S) and PCR-amplifying the region of the gene targeted by each sgRNA. Sequence analysis using Synthego’s Inference of CRISPR Editing (ICE) tool revealed that treatment with one of the sgRNAs (sg2) had induced the highest percentage of cells with genomic deletions in *OSGIN1* from these polyclonal populations (17%), as compared with the other *OSGIN1*-targeting sgRNAs. HeLa cells from this population were then singled-out in 96-well plates in conditioned medium containing 20% FBS (using a BD FACSAria^TM^ cell sorter) and individual clones were allowed to expand before genomic DNA was again extracted, PCR-amplified to sequence part of exon 4 (see oligonucleotide sequences in Table S2) and analyzed with Synthego’s ICE tool. One clonal population that was recovered was shown to have >90% of cells bearing a deletion in *OSGIN1*; this population was considered as mutant for *OSGIN1* and subjected to all analyses. We noticed that overall these cells were somewhat difficult to maintain, as ∼5-10% of cells were found to die with each passage and the percentage of binucleation decreased accordingly. For this reason, all experiments conducted on these *OSGIN1*(-) HeLa cells were done within 7 passages or less, a stage at which ICE analysis indicated that >85% of the clonal population remained mutant for *OSGIN1* and qPCR analysis did not detect the presence of *OSGIN1* transcript. HeLa cells deleted of OSGIN2 were generated following a similar approach, from a polyclonal population with 70% of cells bearing a genomic deletion in exon 2 (targeted by sg4, see oligonucleotide sequences in Table S2). After singling-out in 96-well plates, a monoclonal population was recovered and shown to have >95% of cells bearing a deletion in *OSGIN2*.

The generation of HeLa cells stably expressing the *dTom-2xrGBD* RhoA activity reporter (27) was done by lentiviral transduction as described above, but using pLV-dimericTomato-2xrGBD (a gift from Dorus Gadella; Addgene #176098) as transfer plasmid in HEK293T cells. Supernatants were harvested, spun, filtered, mixed with hexadimethrine bromide (as above) and added to *Rosa26*(-) or *OSGIN1*(-) HeLa cells, with or without stable expression of GFP-HA-OSGN-1. After selection with 1 µg/mL puromycin for 48h, cells single-positive for dTom-2xrGBD or double-positive for GFP and dTom-2xrGBD were sorted using a BD FACSAria^TM^ cell sorter. A polyclonal cell population with minimal *dTom-2xrGBD* expression (displaying a mean fluorescence intensity slightly above the gate of the non-transduced control) was recovered and expanded prior to imaging.

### Immunoblotting of human cell lysates

HeLa cells were plated in 6-well plates (2.5 x 10^5^ cells/well) and grown at 37°C for 48h. Cell were then washed twice on ice with PBS buffer (137 mM NaCl, 2.7 mM KCl, 8 mM Na_2_HPO_4_, and 2 mM KH_2_PO_4_) and harvested by incubating for 30 min at 4°C in 0.6 ml of TGH buffer (50 mM HEPES pH 7.4, 50 mM NaCl, 0.5 mM EDTA, 1% v/v Triton X-100, 10% v/v glycerol). Samples were centrifuged to remove cell debris and an equal volume of 2X Laemmli buffer (65.8 mM Tris pH 6.8, 2.1% SDS, 26.3% (w/v) glycerol, 0.01% bromophenol blue) was added to supernatants. Samples were run on 10% SDS-PAGE and transferred to nitrocellulose. Membranes were blocked for 1h at room temperature in TBST buffer (150 mM NaCl, 50 mM Tris pH 7.6, 0.1% Tween 20) supplemented with 5% powder milk, followed by blotting overnight at 4°C with primary antibodies in TBST buffer. Membranes were washed 3 x 5 min in TBST buffer, incubated at room temperature with horseradish peroxidase-coupled secondary antibodies for 1h in TBST buffer supplemented with 5% powder milk, and washed again 3 x 5 min in TBST buffer. Samples were revealed using Clarity Western ECL Substrate (Biorad, 1705060) and ImageQuant LAS 4000 Imager (GE Healthcare). The following primary antibodies were used: mouse α-α-tubulin (clone DM1A, 1:5000, Sigma, T6199), goat α-GFP (1:1000, Rockland Immunochemicals, 600-101-215), mouse α-6xHis (clone H3, 1:1000, Santa Cruz, sc-8036). The following secondary antibodies were used: goat α-mouse (1:10 000, Bio-Rad, 170-6515), donkey α-goat (1:1000, Invitrogen, A16005).

### RNA isolation and quantitative PCR analysis

HeLa cells were plated in 6-well plates (2.5 x 10^5^ cells/well), grown at 37°C for 48h, washed twice on ice with PBS buffer and lysed by incubating for 5 min at room temperature in 0.6 ml TRIzol Reagent (Invitrogen 15596026). An equal volume of chloroform was added and extracts were centrifuged for 5 min at 4°C. The aqueous phase was collected and an equal volume of isopropanol was added to precipitate RNA. Samples were centrifuged for 10 min at 4°C and the pellets were washed once with ethanol, dried and resuspended in 50 µl ddH_2_O. Synthesis of cDNA was done with 2 µg of total RNA using the High Capacity cDNA Reverse Transcription kit (Applied Biosystems, 4368814). Quantitative PCR (qPCR) was performed in duplicate with 20 ng of cDNA per reaction using the Taqman Fast qPCR MasterMix (Applied Biosystems, 4444557) and reactions were run for 40 cycles on a QuantStudio™ 7 Flex Real-Time PCR System, using primer sets specific for *OSGIN1* and *OSGIN2* and from the Universal ProbeLibrary (Roche, see Table S2 for sequences). Analysis was done with Expression Suite software (Applied biosystems) using the genes encoding ß-actin (*ACTB*) and GAPD (*GAPDH*) as controls.

### Time-lapse confocal microcopy

*C. elegans* embryos were obtained by cutting open synchronized gravid hermaphrodites in egg buffer (25 mM HEPES pH 7.3, 118 mM NaCl, 48 mM KCl, 2 mM CaCl_2_, 2 mM MgCl_2_) using two 25-gauge needles and individually mounted on a coverslip. The coverslip was flipped on a 3% agarose pad on a glass slide and sealed with valap (1:1:1 vaseline:lanolin:paraffin). For RNAi-depleted animals, time-lapse images were acquired with an Apo LWD 40x/1.25 NA water-immersion objective mounted on a Nikon A1R laser-scanning confocal microscope, and 0.8 µm-thick confocal sections of the entire embryo were obtained by illumination in single-track mode with 488 nm and 561 nm laser lines controlled by NIS Elements 4.2 software (Nikon). For *osgn-1* mutant animals, time-lapse images were acquired with a Plan-Apocromat 63x/1.4 NA oil-immersion objective mounted on a Zeiss LSM700 laser-scanning confocal microscope, and 0.8 µm-thick confocal sections of the entire embryo were obtained by illumination in single-track mode with 488 nm and 555 nm laser lines controlled by ZEN 2011 software (Zeiss). In both cases, images were acquired every 3 min for the first 45 min, and then every 15 min for 2 hr. Anaphase onset (t = 0 min) was defined as the first frame where two sets of chromosomes are visible.

HeLa cells were plated on 35 mm microscopy dishes (MatTek, P35G-1.5-14-c) and left to adhere overnight before co-transfection (using Lipofectamine 3000 treatment, as above) of vectors enabling expression of H2B-mRFP and rGFP-CAAX, or mRFP-CAAX alone. After 48h, cells in prometaphase or metaphase were selected from asynchronous populations. For measurements of cytokinesis progress and membrane stability, time-lapse images were acquired for 10h at 5-min intervals with a Plan-Apocromat 40x/1.4 NA oil-immersion objective mounted on a Zeiss LSM700 laser-scanning confocal microscope controlled by Zen software, acquiring 0.8 µm-thick confocal sections of the entire cell using single-track mode illumination with 488 and 555 nm laser lines. For visualization of the dTom-2xrGBD reporter, time-lapse images were acquired for 18h at 3-min intervals with a HC PL APO 63X/1.4 N.A oil-immersion objective mounted on a Leica SP8 laser-scanning confocal microscope controlled by LAS X software (Leica), using single-track mode illumination with 488 and 552 nm laser lines. Anaphase onset (t = 0 min) was again defined as the first frame where two sets of chromosomes are visible.

### Quantitative image analysis

All images were processed and analyzed from the original files using ImageJ software (National Institute of Health) and are represented as sum or maximal intensity projections converted into black and white renderings (see also figures legends). For better visualisation, brightness and contrast were adjusted using the same settings for all images within a panel.

For measurements of GFP::AHPH reporter levels at the equator of dividing P_4_ embryonic blastomeres, fluorescence integrated density was acquired throughout the entire movie in 5 successive confocal planes within a 8 μm x 3 μm region of interest (ROI) positioned at the cell equator. For each image, fluorescence integrated density was also measured in two regions of 2 μm x 6 μm located within the cytoplasm of each daughter cell. After bleach correction (from a curve generated by measuring the entire imaging field), the sum of values obtained within the cytoplasm were subtracted from the sum of values obtained from the cell equator at each time point. Position of the P_4_ blastomere can vary within the confocal plane, and therefore measurements were made on embryos in which P_4_ was found within 15 confocal sections of the coverslip. GFP::AHPH reporter levels at the intercellular bridge of somatic blastomeres was measured in the same manner in neighboring cells that divided at the same time as the P_4_ blastomere.

For measurements of dTom-2xrGBD reporter levels at the cytokinetic furrow and midbody of dividing HeLa cells, fluorescence integrated density was acquired throughout the entire movie in a single confocal plane along a 10 pixel-thick line drawn across the cell equator. Integrated density was also measured along 2 lines drawn on each side of the equator, passing through the cytoplasm of each daughter cell. After bleach correction (from a curve generated by measuring the entire imaging field), enrichment of the reporter was expressed as a ratio of the area under the curve (AUC) of cell cortices measured at the equator (5 pixels/cortex) to that of cell cortices measured on each side, in each daughter cell.

Cytokinetic furrow ingression rate and intercellular bridge stability were scored by tracking plasma membrane signal, in both HeLa cells (using rGFP-CAAX or mRFP-CAAX probes) and in *C. elegans* embryos (using the mCherry::PH^PLC61^ probe), throughout the entire movie. Formation of the intercellular bridge was observed as a progressive coalescence of fluorescence peaks along a line drawn at the cell equator, and bridge collapse was defined as a subsequent separation of this single peak into two distinct peaks.

### Indirect immunofluorescence

HeLa cells were grown as asynchronous populations on 6-well imaging plates (1.5 x 10^5^ cells/well) for 24h prior to transfection (using Lipofectamine 3000 treatment, as above). Media was aspirated 48h later and cells were fixed either by treatment for 5 min at room temperature (RT) with paraformaldehyde (4% v/v in PBS buffer; for anti-GFP or anti-HA staining), or by treatment for 10 min on ice with trichloroacetic acid (10% v/v in PBS buffer supplemented with 20 mM glycine (G-PBS); for anti-RhoA staining). Fixed cells were washed three times in PBS buffer (or G-PBS for anti-RhoA staining), permeabilized for 5 min at RT in 0.3 % Triton X-100 in PBS buffer (or 0.2% Triton X-100 in G-PBS buffer for anti-RhoA staining), and blocked for 20 min at RT in PBS (or G-PBS) buffer supplemented with 2% bovine serum albumin (BSA). The cells were then incubated for 1h at RT with primary antibodies diluted in PBS (or G-PBS) buffer supplemented with 0.5% BSA, washed 3 times with PBS (or G-PBS) buffer, and incubated for 1h at RT with Alexa fluor dye-conjugated secondary antibodies (1:500) and Alexa-555-coupled phalloidin (1:400, Invitrogen, A34055) diluted in PBS buffer supplemented with 0.5% BSA. After three washes with PBS buffer, cells were then treated for 5 min at RT with Hoechst 33342 (1 µg/mL in PBS buffer, Invitrogen, H1399), followed by two additional washes of 10 min each with PBS buffer. For co-staining with goat α-GFP antibody, secondary antibody was added prior to addition of other primary antibodies to diminish background due to nonspecific interactions. Coverslips were mounted/sealed on glass slides using Prolong Gold Antifade Reagent (Invitrogen, P10144) and allowed to dry overnight at RT before imaging. Images were acquired with a Plan-Apocromat 40×/1.4 NA oil-immersion objective mounted on a Zeiss LSM700 laser-scanning confocal microscope, and 0.8 µm-thick confocal sections of the entire cells obtained by illumination in multi-track mode with 405, 488 and 555 nm laser lines controlled by Zen software (Zeiss). The following primary antibodies were used: goat α-GFP (1:1000, Rockland Immunochemicals, 600-101-215), mouse α-HA (clone 12CA5, 1:200, Sigma, 11583816001), rabbit α-pMLC-S20 (1:1000, Abcam, AB2480), rabbit α-RhoA (clone 67B9, Cell Signaling Technology, 2117T). The following Alexa Fluor-coupled secondary antibodies were used: donkey

α-rabbit (Invitrogen, A10042), donkey α-mouse (Invitrogen, A31571), donkey α-goat (Invitrogen, A11055).

## Statistical analysis

GraphPad Prism software was used for graphing and all statistical analyses. Parametric tests of statistical significance between samples were applied except when assumptions of normality and equal variance were not met, see figures legends for details. In all cases a two-tailed *p*-value smaller than 0.05 was considered significant. All results are expressed as average ± standard deviation (SD) or standard error of the mean (SEM). Sample size (*n*) is given in each figure panel or legend. All results shown are representative of at least three independent biological replicates (N) for each condition.

## Author contributions

E.G., L.L. and J.-C.L. designed the experiments. E.G. performed experiments and analyzed data for all figures except Fig. 3F. L.L. generated OSGN-1 mutant constructs, performed experiments and analyzed data for Fig. 3 and 4. J.B. and S.G. contributed to the phenotypic analysis of *C. elegans* mutants. M.K.S.E.L. and S.M. designed the sgRNA sequences and helped to generate the HeLa cells deleted of *OSGIN1* and *OSGIN2*. E.G. and J.-C.L. wrote the manuscript, with editing input from all authors.

## Acknowledgements

We thank Sébastien Carreno, Audrey Claing, Étienne Gagnon, Michael Glotzer, Alisa Piekny, Guy Sauvageau and Mike Tyers for strains, constructs, cell lines and reagents. We also thank Joaquim Javary for help with viral transductions and Almer van der Sloot for help with OSGN-1 structural analysis. We are also grateful to Christian Charbonneau, Annie Gosselin, Angélique Bellemare-Pelletier and Raphaëlle Lambert of the Institute for Research in Immunology and Cancer’s (IRIC’s) Bio-imaging, Flow cytometry and Genomic Facilities for technical assistance, Rébecca Panès and Angélique Bellemare-Pelletier for lentiviral production, Victor Poulin-Therrien for help with *C. elegans* strains outcrossing, Abby Gerhold and Greg FitzHarris for critical reading of the manuscript, and all members of the FitzHarris, Gerhold, Hickson and Labbé laboratories for helpful discussions. Some strains were provided by the CGC, which is funded by the NIH Office of Research Infrastructure Programs (P40 OD010440). L.L. is a recipient of studentships from the Fonds de la Recherche du Québec - Santé (FRQ-S), the Canadian Institutes of Health Research (CIHR), Université de Montréal’s Graduate Studies and the IRIC. This study was supported by grants from the CIHR (PJT-480641) and the Cancer Research Society (24475) to J.-C. L.

## Declaration of interests

The authors declare no competing interests.

## Supplemental Information

**Table S1.**
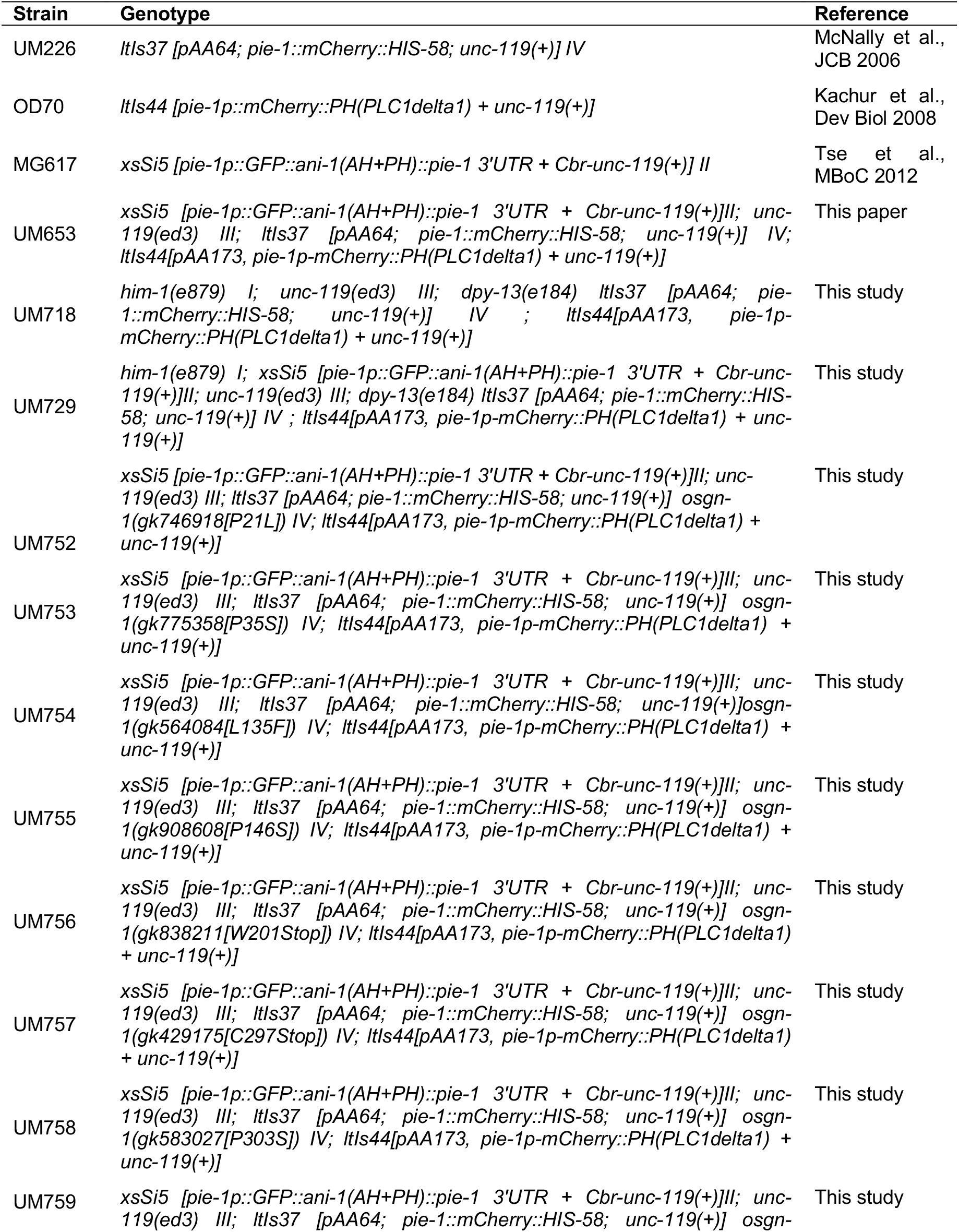
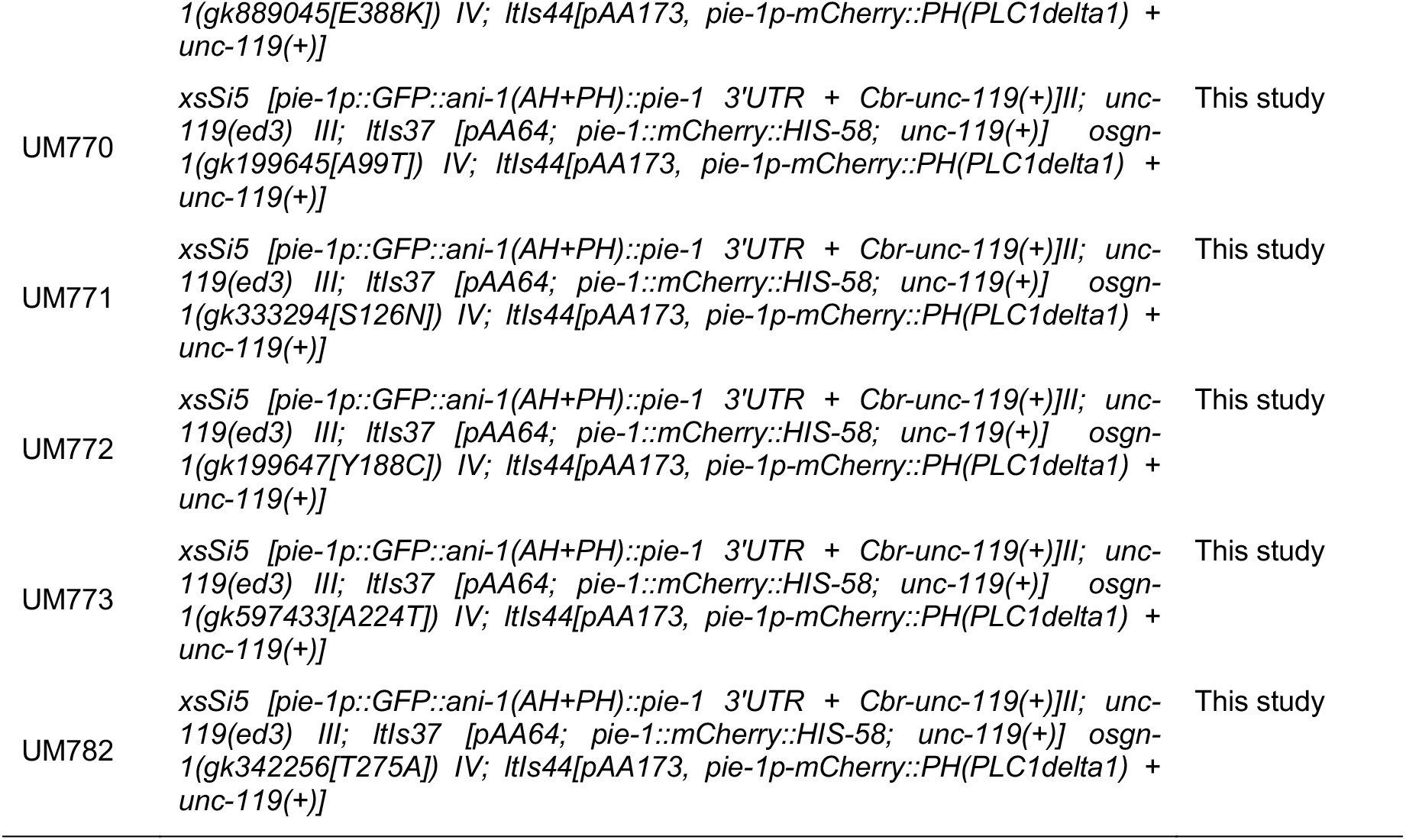
*C. elegans* strains.

**Table S2.**
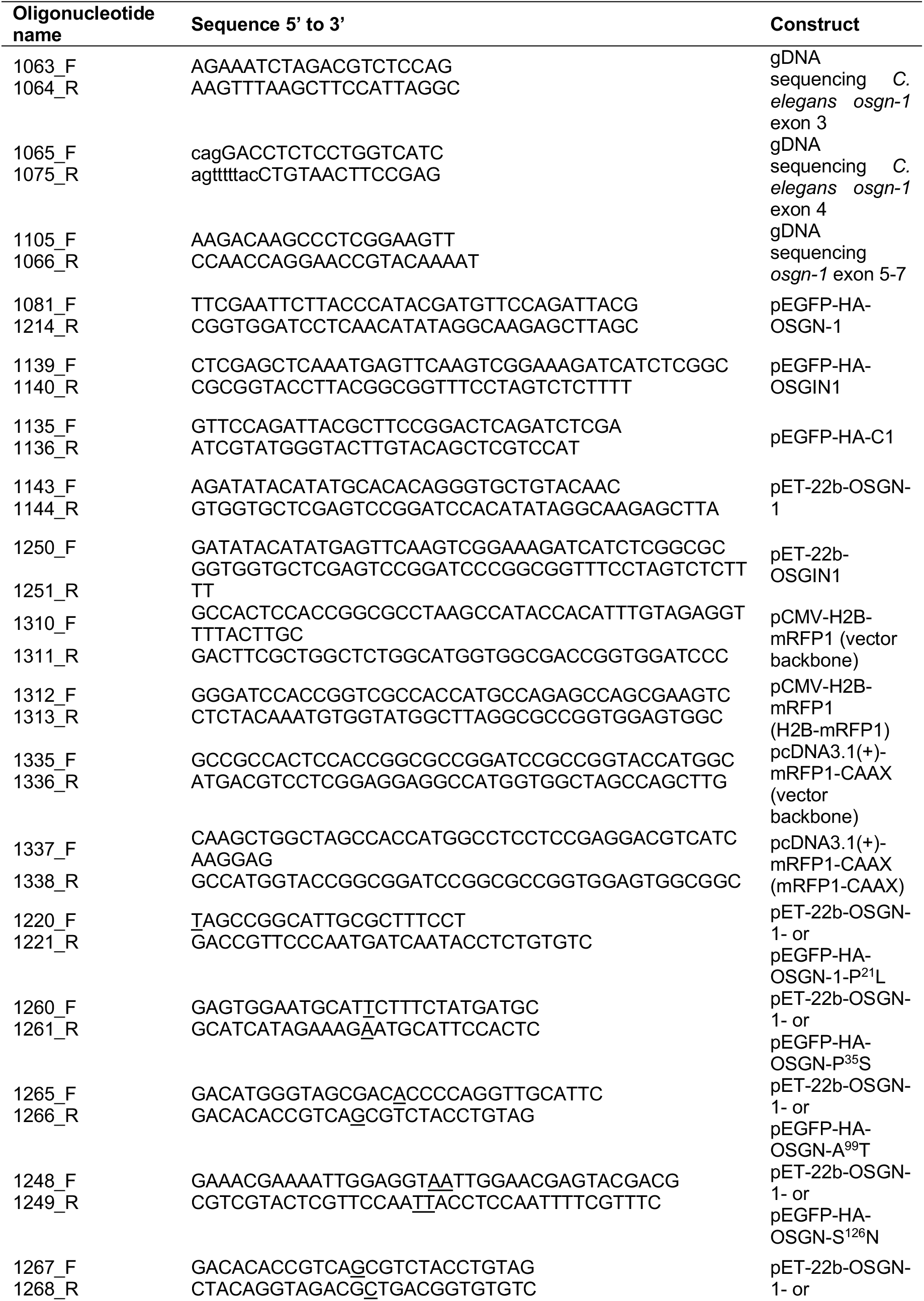
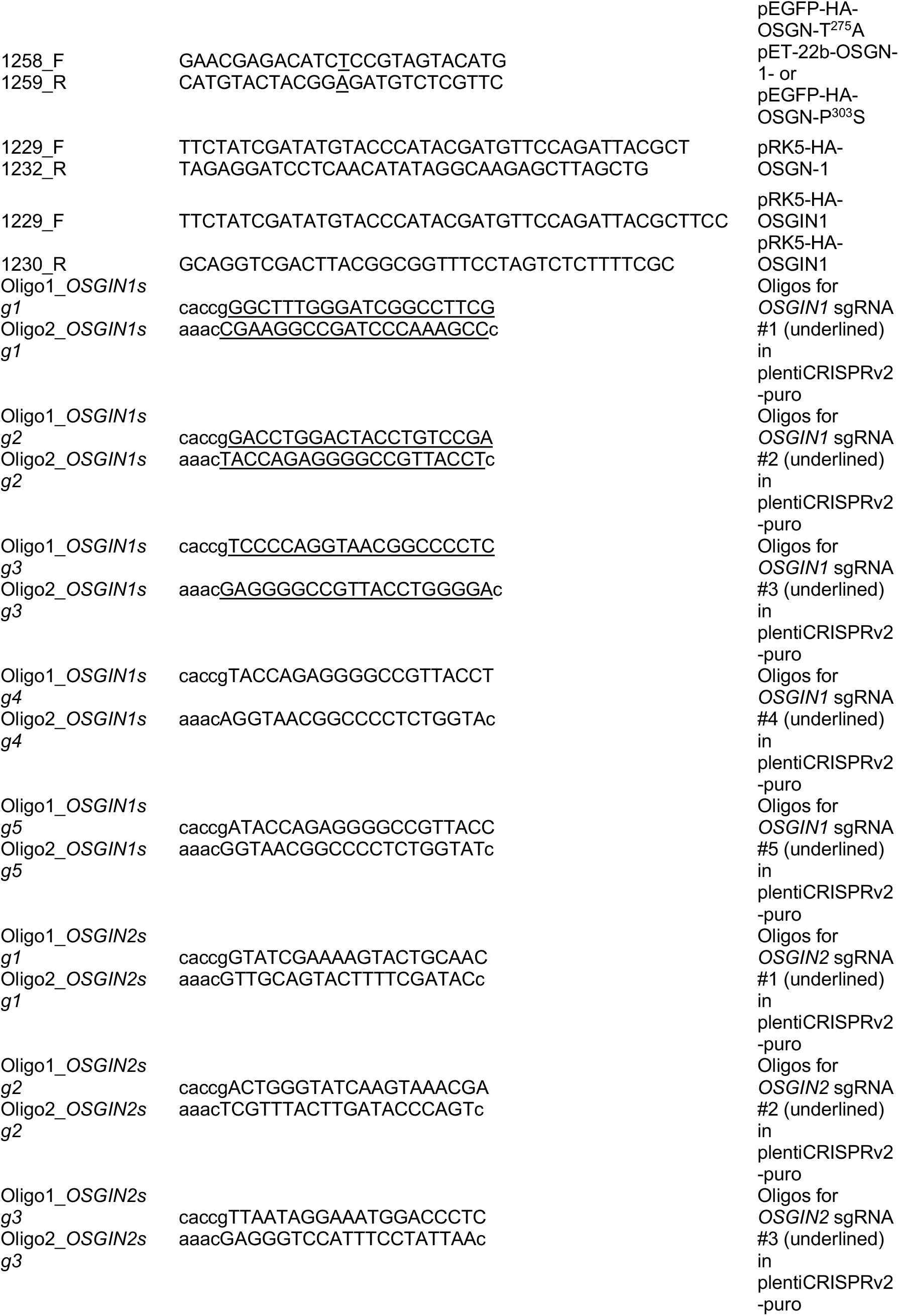
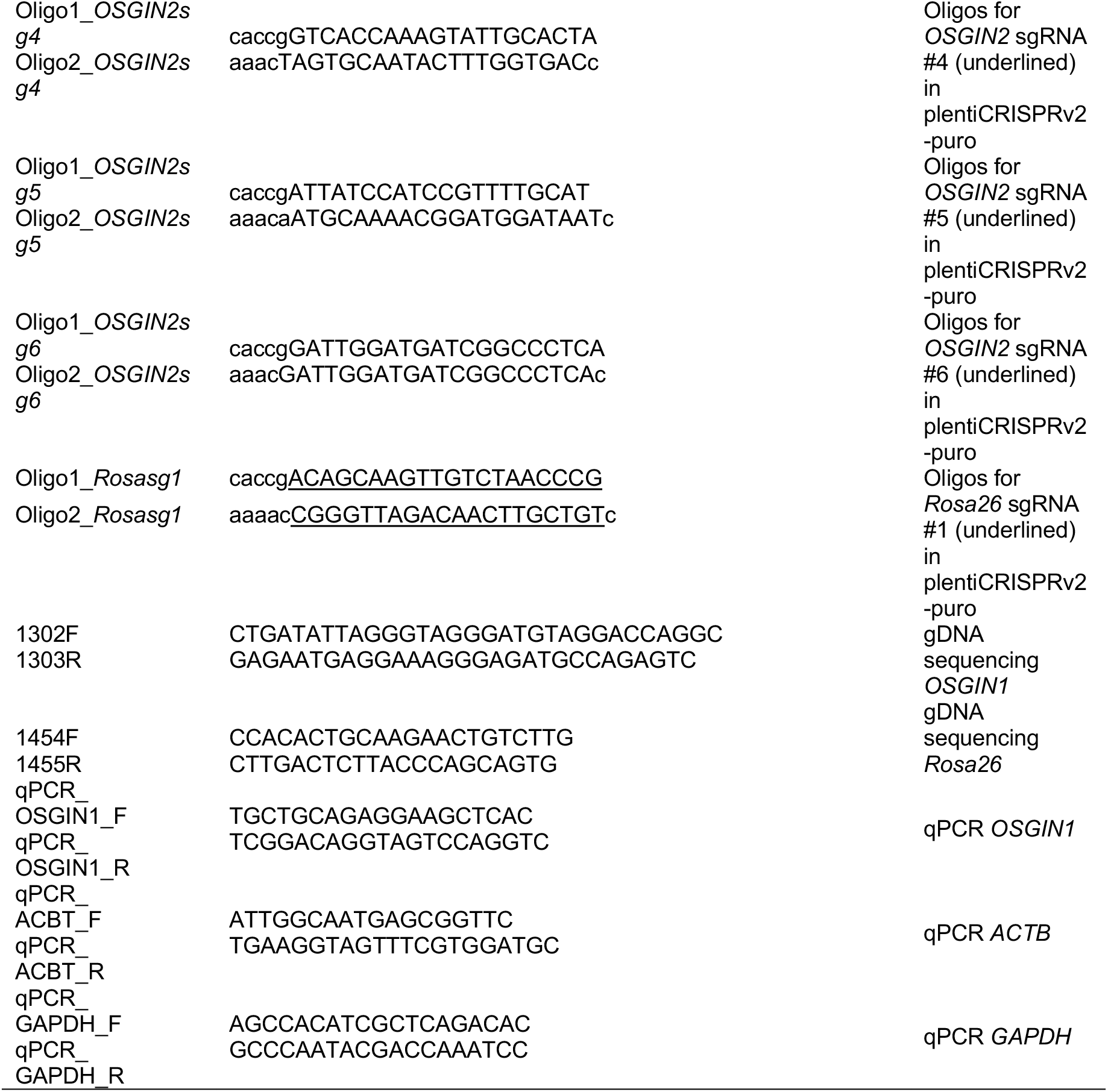
Oligonucleotide sequences.

**Table S3.**
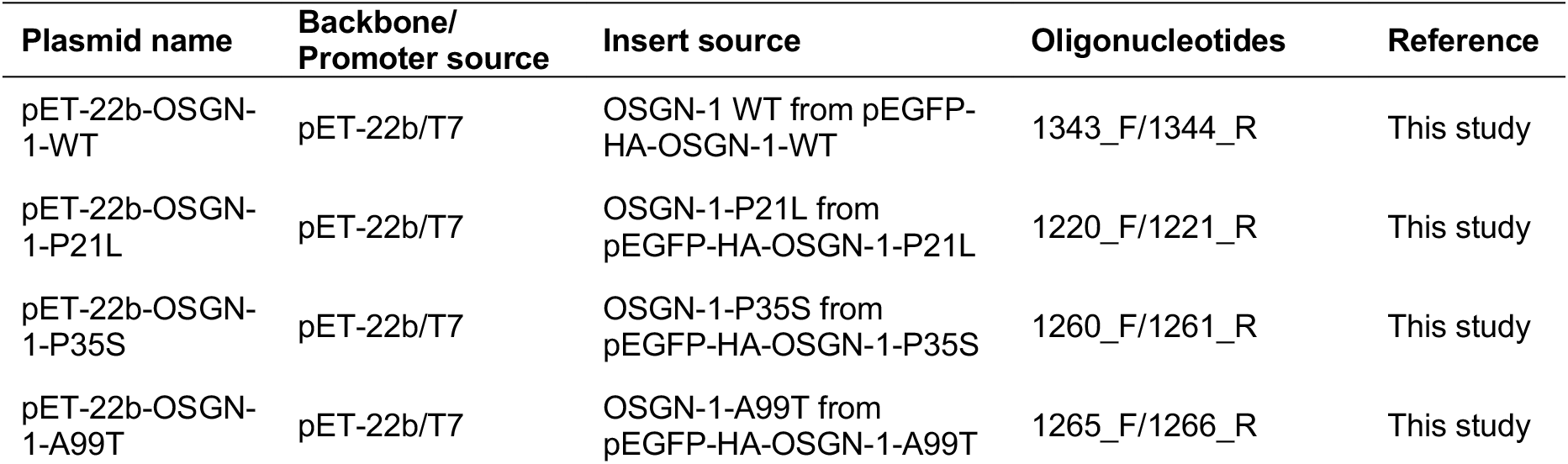
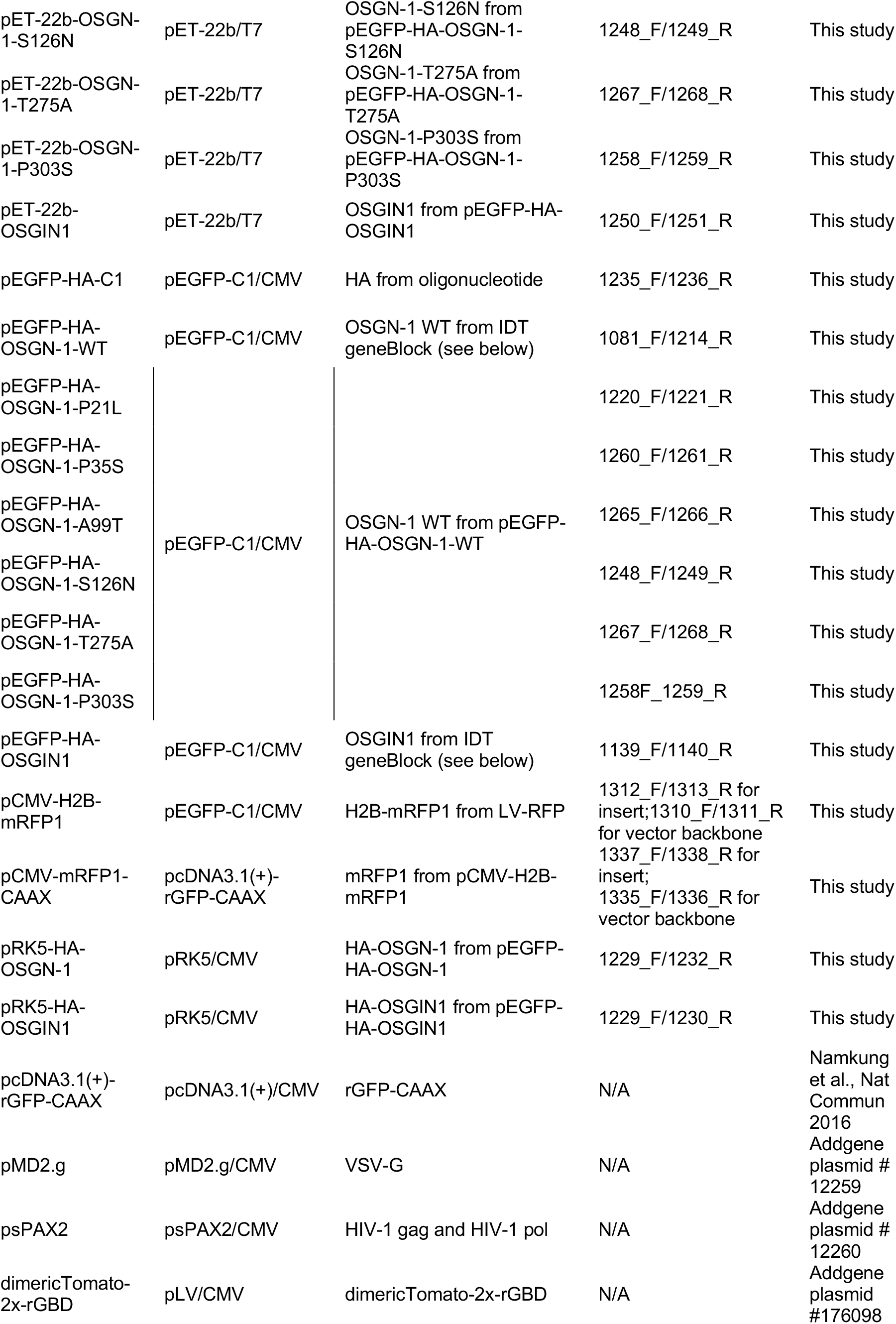
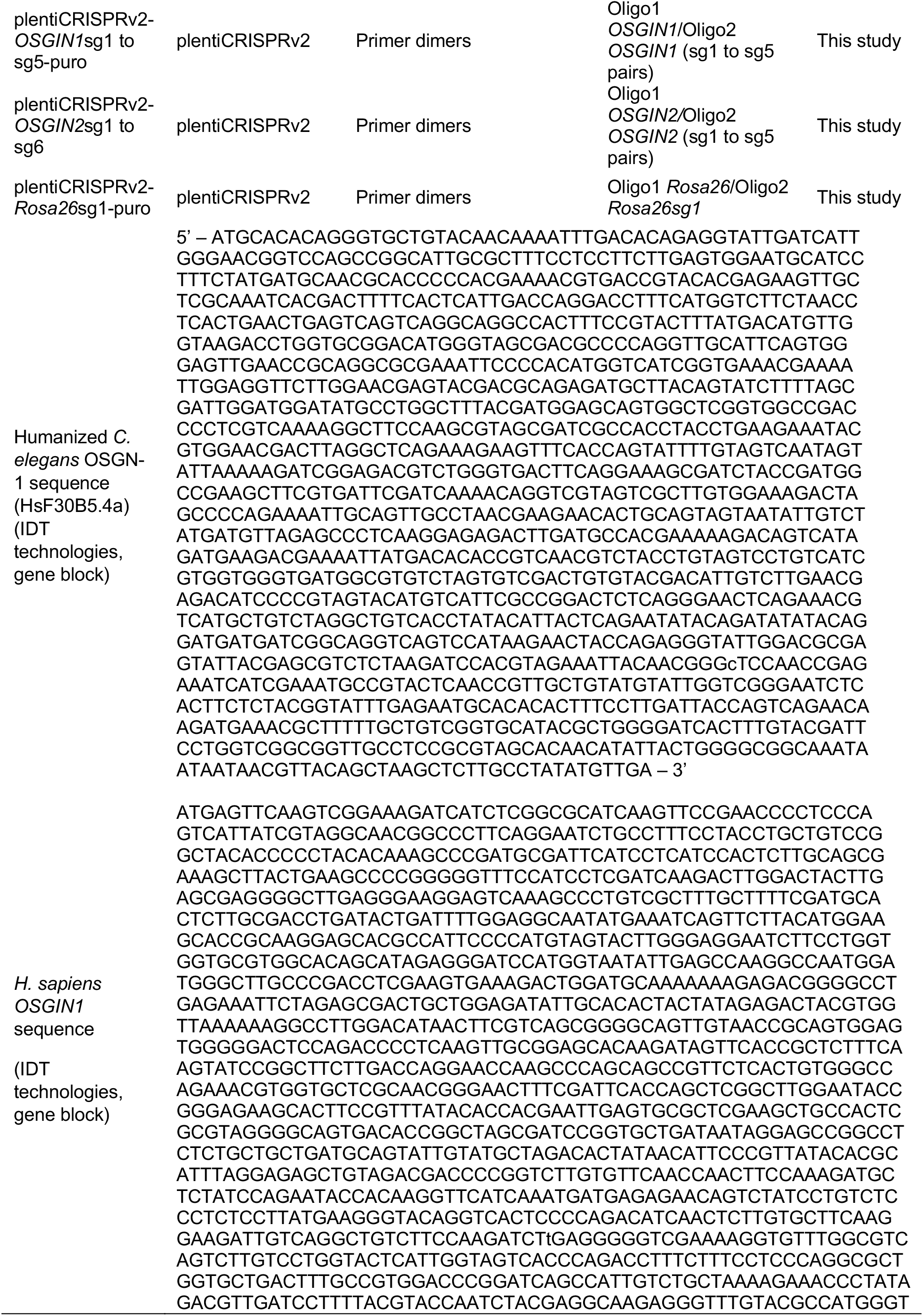
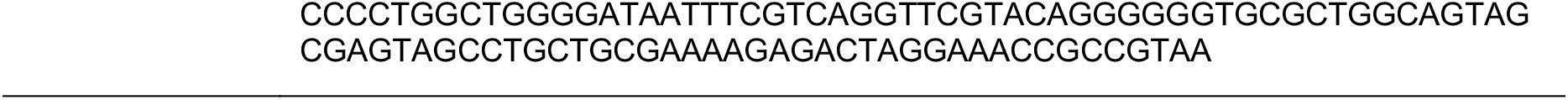
DNA constructs.

## Supplemental Figures

**Figure S1.**
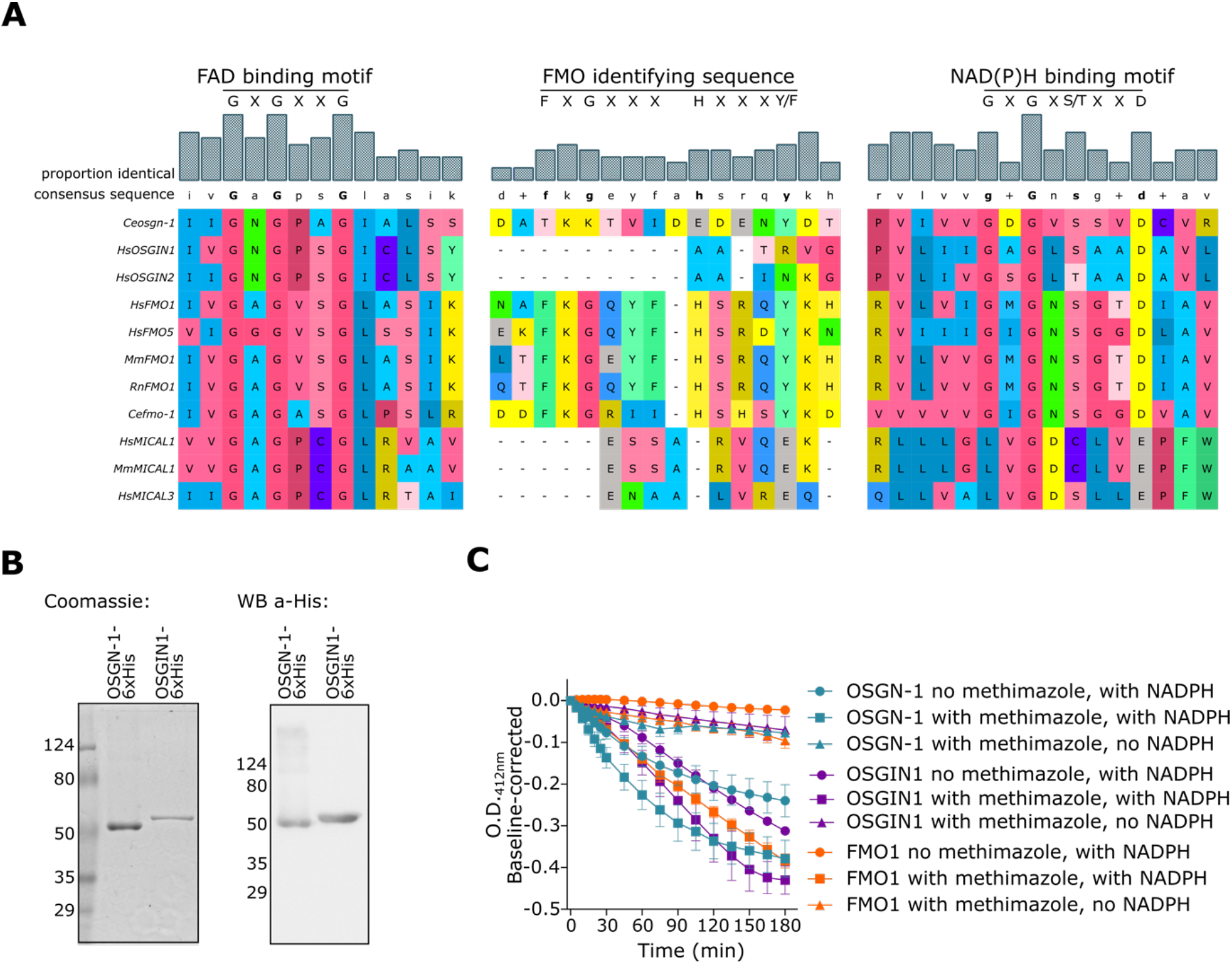
*C. elegans* OSGN-1 and human OSGIN1 are flavin-containing monooxygenase enzymes. (A) Sequence alignments of the FAD, MO and NADPH domains of *C. elegans* OSGN-1 and human OSGIN1 and OSGIN2 with those of canonical and non-canonical FMOs, namely human, mouse, rat and *C. elegans* FMO1, human FMO5, human and mouse MICAL1, and human MICAL3. (B) Coomassie straining (left) and Western blot analysis revealed with anti-His antibodies (right) of bacterially-purified *C. elegans* OSGN-1-6xHis and human OSGIN1-6xHis. (C) Comparative measurement of the catalytic activity of human microsomal FMO1 (orange), *C. elegans* OSGN-1-6xHis (cyan) and human OSGIN1-6xHis (magenta; 2 μM final concentration each) using the DTNB-methimazole assay, in the presence of both methimazole and NADPH (squares) or in absence of methimazole (circles) or NADPH (triangles). The decrease of absorbance at 412 nm over time (in min) reports on catalytic activity. Values are corrected for the initial measurement at the start of the reaction and correspond to the mean ± SEM over N = 3 assays.

**Figure S2.**
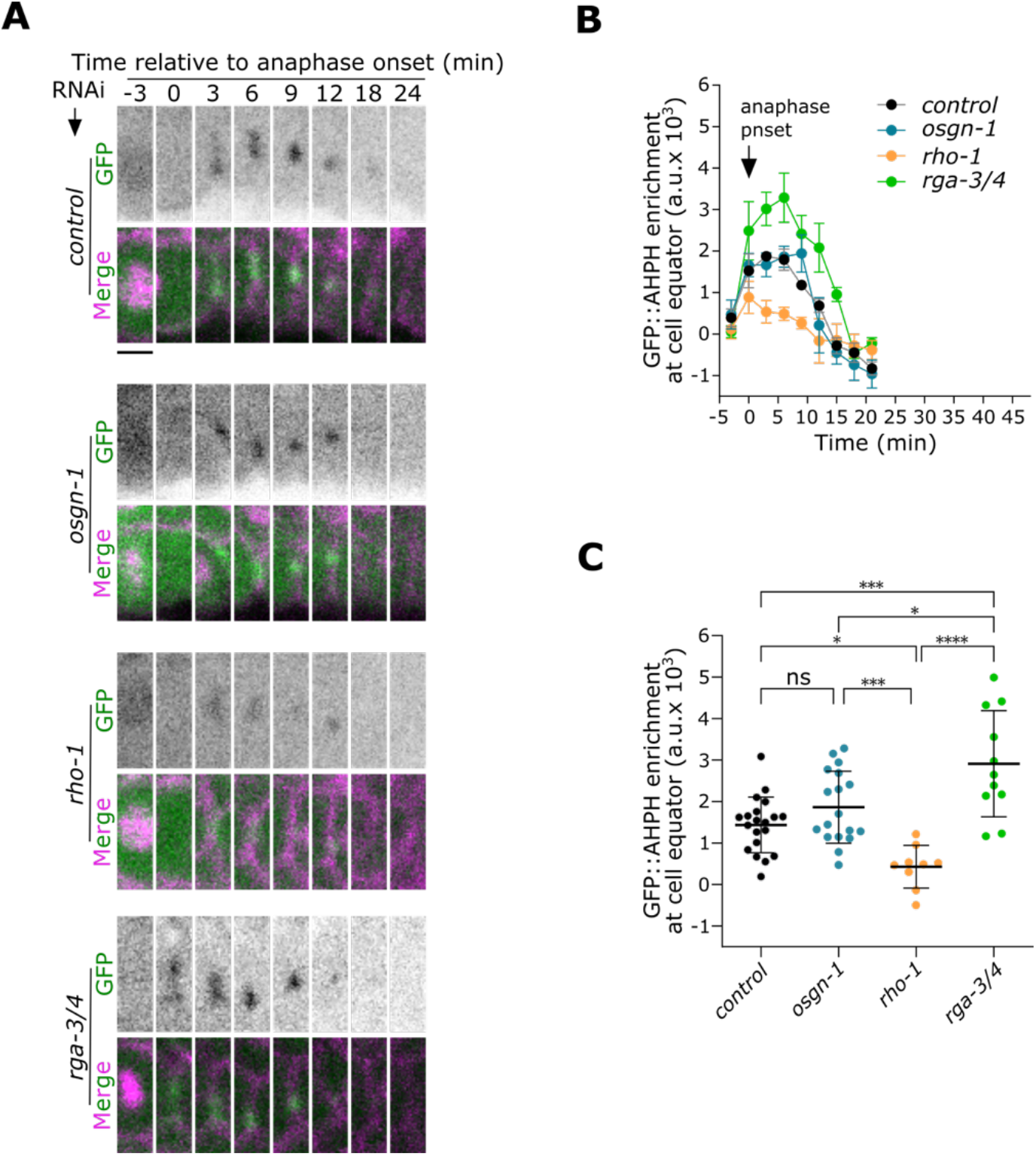
OSGN-1 does not impact RhoA activity in embryonic somatic cells. (A) Time-lapse images (sum projections of three confocal slices) of somatic cell divisions in the same *C. elegans* embryos represented in Fig. 2 and depleted of control (empty L4440 vector), *osgn-1*, *rho-1* or *rga-3/4* by RNAi. Embryos express markers for RhoA activity (GFP::AHPH, green), membrane (mCh::PLC∂-PH, magenta) and chromatin (mCh::H2B, magenta). Time (in min) is relative to anaphase onset (A.O.). Scale bar = 3 μm. (B) Fluorescence levels over time of the GFP::AHPH reporter measured at the equatorial region (cytokinetic furrow/intercellular bridge) of embryos depleted of control (black), *osgn-1* (blue), *rho-1* (orange) or *rga-3/4* (green) by RNAi, as in panel a. Values correspond to mean ± SEM over 3 biological replicates (n=9-20 embryos in total). (C) Average fluorescence intensity of the GFP::AHPH reporter measured in somatic cells during early cytokinetic furrow ingression (6-9 min after A.O.) in embryos depleted of the indicated gene products. Values represent individual embryos. Bars denote average ± SD (n=9-20 embryos). ns = non-significant, *p < 0.05, ***p < 0.001, two-way ANOVA followed by a Šídák multiple comparison test.

**Figure S3.**
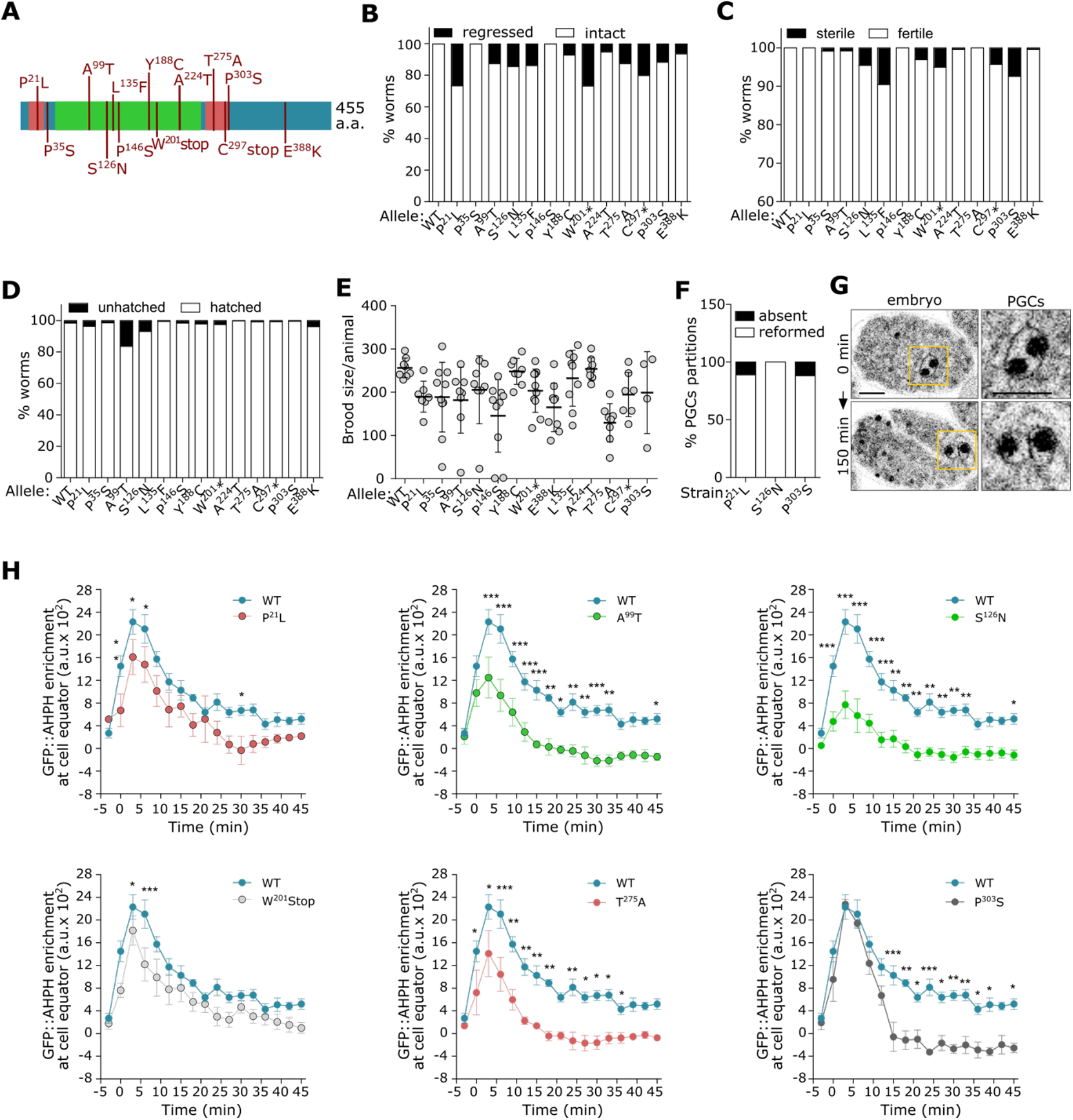
Phenotypic characterization of *C. elegans osgn-1* mutants. (A) Schematic depiction of the different amino acid substitutions encoded by the *osgn-1* mutants used in this study. (B-D) Quantification of the percentage of PGC bridge stability (B), fertility (C) and embryonic viability (D) in animals bearing the indicated *osgn-1* allele. (E) Quantification of the total brood size per animal bearing the indicated *osgn-1* allele. Values represent individual animals. Bars denote average ± SD (n=99-372 animals). (F-G) Quantification (F) and representative images (G) of the presence (white bars) or absence (black bars) of a membrane partition between the two PGC nuclei in late embryos bearing the indicated *osgn-*1 allele and expressing markers for RhoA activity (not shown), membrane (mCh::PLC∂-PH, black) and chromatin (mCh::H2B, black). All depicted embryos had previously been visually documented to have undergone regression of the membrane between the two PGCs. The membrane and nuclei markers are shown. Scale bar = 10 μm, n = 8-17 embryos. (H) Fluorescence levels over time of the GFP::AHPH reporter measured at the equatorial region (cytokinetic furrow/intercellular bridge) of P_4_ in control embryos (black) or in embryos bearing the indicated *osgn-*1 allele (colored), as in Fig. 3. In each graph time is relative to anaphase onset (A.O.). Values correspond to mean ± SEM over 3 biological replicates (n=13-23 embryos each in total). Ns = non significant, *p < 0.05, **p < 0.01, ***p < 0.001, two-way ANOVA with repeated measures followed by a Dunnett multiple comparison test.

**Figure S4.**
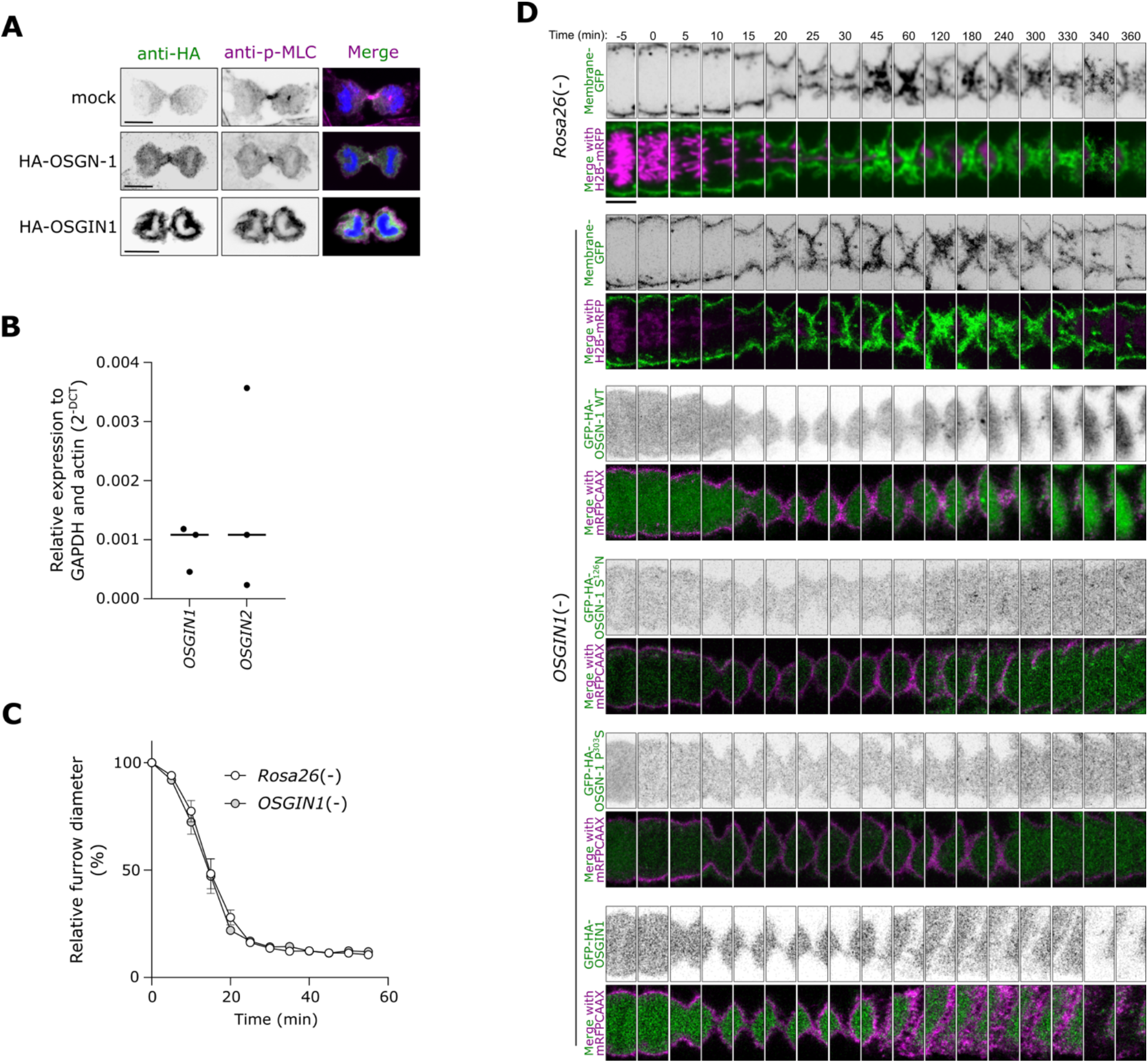
*C. elegans* OSGN-1 is the functional ortholog of human OSGIN1. (A) Representative images of indirect immunofluorescence against HA (green) and p-Myosin Light Chain (p-MLC, magenta), in asynchronous HeLa cells transfected with an empty vector (mock), HA-OSGN-1 or HA-OSGIN1. Hoechst labels the nuclei (blue). Scale bar = 10 μm. (B) Quantitative PCR analysis of *OSGIN1* and *OSGIN2* mRNA levels in HeLa cells in three independent biological replicates. (C) Cytokinetic furrow diameter over time (in min) relative to the diameter at the start of anaphase (time 0) in dividing HeLa cells deleted of *OSGIN1* or *Rosa26* (from Fig. 4f). (D) Time-lapse images of cytokinetic furrow ingression in *Rosa26* control or *OSGIN1* mutant HeLa cells co-expressing either markers for the plasma membrane (GFP-CAAX, green) and nuclei (H2B-mRFP, magenta), or a marker for the plasma membrane (mRFP-CAAX, magenta) and a GFP-tagged *C. elegans* OSGN-1 variant (WT or bearing the S^126^N or P^303^S mutations) or human OSGIN1, as indicated. The membrane undergoes regression in cells that express S^126^N or P^303^S variants of OSGN-1. Scale bar = 10 μm.

## Notes

### Competing Interest Statement

The authors have declared no competing interest.

